# Interacting abiotic and biotic drivers shape woody invasions across Hawaiian forests

**DOI:** 10.1101/2022.06.01.494375

**Authors:** Dylan Craven, Jonathan M. Chase, Tiffany M. Knight

**Affiliations:** Centro de Modelación y Monitoreo de Ecosistemas, Facultad de Ciencias, Universidad Mayor, Santiago, Chile; German Centre for Integrative Biodiversity Research (iDiv) Halle-Jena-Leipzig, 04103 Leipzig, Germany; Department of Community Ecology, Helmholtz Centre for Environmental Research–UFZ, 06120 Halle (Saale), Germany; Department of Computer Science, Martin Luther University, Halle-Wittenberg, 06108 Halle (Saale), Germany; Institute of Biology, Martin Luther University Halle-Wittenberg, Am Kirchtor 1, 06108 Halle (Saale), Germany

**Keywords:** biological invasions, context dependency, oceanic islands, Darwin’s Naturalisation Hypothesis, functional traits

## Abstract

The same features that generate native biodiversity patterns across and within oceanic islands over evolutionary time – climate, soil age, topography, and biotic interactions – also influence their vulnerability to biological invasions. Here, we identify the factors that shape the richness and abundance of alien woody species in forest communities across the Hawaiian archipelago, and assess the relative importance of abiotic, biotic, and anthropogenic factors and their interactions on the establishment and dominance of woody alien species. Using a database of 460 forest plots distributed across the six major Hawaiian islands, we examine variation in *i*) relative alien species richness and abundance as a function of abiotic and anthropogenic factors (e.g., temperature, aridity, soil age, and the human influence index) and *ii*) establishment and dominance of alien species as a function of the same abiotic and anthropogenic factors, as well as phylogenetic and trait distinctiveness. We found that relative alien species richness and abundance were higher in areas where temperature was high and aridity low. Gradients in temperature, aridity, soil age, and human influence also modulated the importance of biotic factors in determining establishment of alien species. In contrast, whether these alien species could become locally dominant was not strongly influenced by abiotic or biotic factors, or their interactions. Our results suggest that environmental filtering mediates the strength of biotic filtering in determining where woody aliens are able to colonize and establish on these oceanic islands, but not whether they become dominant. The context dependence of multi-species invasions highlights the complexity of developing management strategies to mitigate the biodiversity and ecosystem impacts of biological invasions.

## Introduction

On just 3 to 5% of the world’s area, islands harbor a disproportionate amount (15% to 20%) of its known plant diversity (Tershy et al. 2015, Whittaker et al. 2017). Especially on oceanic islands, this biodiversity has emerged as a result of isolation and dynamic geophysical processes that influence island area and environmental heterogeneity (Gillespie 2016, Schluter and Pennell 2017, Whittaker et al. 2017). However, these isolated islands are also highly susceptible to invasions by alien species (Moser et al. 2018), which can alter community structure, displace endemic species, and threaten the provisioning of vital ecosystem services (Vitousek et al. 1987, Sax and Gaines 2008, Vilà et al. 2011, Simberloff et al. 2013, Castro-Díez et al. 2019). There is no sign that the number of alien species that are successfully establishing is reaching equilibrium (Seebens et al. 2017), suggesting that without prevention and management, new alien species will continue to arrive and disrupt island ecosystems. It is therefore critical to deepen current understanding of the factors that allow alien species to successfully establish and come to dominate in island ecosystems (Mack et al. 2000, Pyšek et al. 2010, Blackburn et al. 2011, Dawson et al. 2017).

Across and within oceanic islands, abiotic, biogeographical, and anthropogenic factors jointly determine where biological invasions occur. Alien plant floras are typically more diverse on islands that are isolated, large, and topographically complex, as well as on islands with high human impacts (Sax and Gaines 2008, Denslow et al. 2009, Dawson et al. 2017, Moser et al. 2018, Wohlwend et al. 2021). Within islands, richness and abundance of alien plants tend to be greater at lower elevations in locations with high precipitation (Ibanez et al. 2019) and where human population density is higher and land use is more intense (Denslow 2003). However, community-level patterns of biological invasions often fail to provide mechanistic insights, in part because they overlook functional and phylogenetic differences between native and alien species (biotic filtering). A growing body of literature has emerged to fill this gap, which also highlights that biological invasions are multi-faceted processes most appropriately addressed by jointly considering multiple potential mechanisms, including environmental and biotic filtering, propagule pressure, and human impacts (Catford et al. 2009, Gallien and Carboni 2017, Pearson et al. 2018, Bennett 2019).

Environmental filtering structures communities by influencing the establishment of species in a given place via their traits that determine tolerance of abiotic factors (e.g. water availability, soil pH) (Kraft et al. 2015). This environmental filtering can result in the co-occurrence of species with similar traits. In the context of biological invasions, environmental filtering is frequently framed in terms of the pre-adaptation hypothesis (Cadotte et al. 2018, Bennett 2019), which posits that alien species should be sufficiently similar to native species in their traits or evolutionary history to allow them to establish viable populations. Support for this hypothesis is particularly expected in environments with high abiotic stress such as those with high aridity, cold temperatures, or young soils (Cornwell and Ackerly 2009). Yet empty niche space on islands, created by the rarity and non-randomness of colonizations (Losos and Ricklefs 2009, Carvajal-Endara et al. 2017, Taylor et al. 2019), may occur even in environments with high abiotic stress. For example, Henn et al. (2019) showed that traits of alien plant species were more different from those of native plant species in warm, arid environments on the lower slopes of Mauna Loa, Hawai’i compared to those in cooler, wetter environments, suggesting that more unfilled niche space was available in environments with greater abiotic stress. Empirical evidence in support of the preadaptation hypothesis on island ecosystems (and in general) is therefore quite mixed (Schaefer et al. 2011, Marx et al. 2016, Ma et al. 2016, Cadotte et al. 2018).

On the other hand, biotic filtering can promote the co-occurrence of ecologically dissimilar species by minimizing competition (Kraft et al. 2008, Kraft and Ackerly 2010). However, these species may still not completely fill available niche space (Ashby et al. 2017). For example, dispersal limitation might result in the absence of particular trait syndromes on oceanic islands (Taylor et al. 2019, König et al. 2021). This leaves niche space available for colonization by radiating lineages, such as lobeliads in the Hawaiian Islands (Asterales: Campanulaceae; Givnish et al. 2009) and the *Periploca laevigata* complex in the Canary Islands (García-Verdugo et al. 2020), or by alien species. Assuming that alien species can persist in the new environment, Darwin’s naturalisation conundrum (Darwin 1859) predicts that alien species are more likely to establish in native communities in which they are sufficiently distinct from native species, either phylogenetically or functionally, to minimize competition by exploiting unused niche space (Cadotte et al. 2018, Bennett 2019). A meta-analysis using phylogenetic relatedness as a surrogate for functional differences found that biotic filtering influences the establishment of alien species at local spatial scales in a manner consistent with Darwin’s naturalisation hypothesis (Ma et al. 2016). While phylogenetic dissimilarity is a reliable proxy for detecting differences in how species occupy multidimensional niche space (Tucker et al. 2018), differences in traits associated with competition and resource use (Kunstler et al. 2012, Díaz et al. 2016) provide a more direct means for understanding the relationship between ecological similarity and invasion (Marx et al. 2016, Carboni et al. 2018, van der Sande et al. 2020, Levin et al. 2020).

There is a growing appreciation that biological invasions are context dependent, as they are frequently the result of interactions among abiotic, biotic, and anthropogenic factors (Ricciardi et al. 2013, Pearson et al. 2018, Sapsford et al. 2020, Catford et al. 2022). Along abiotic gradients, native communities are expected to be less invaded and alien plant species more similar to native communities where abiotic stress is high, while native communities are expected to be more invaded and alien species less similar to native communities where abiotic stress is low (Perelman et al. 2007, Tecco et al. 2010, Gross et al. 2013, González-Moreno et al. 2014, Sapsford et al. 2020). Anthropogenic disturbances (or habitats) may also generate context dependent invasion outcomes (González-Moreno et al. 2014, Jauni et al. 2015) by favoring non-native species with disturbance-tolerant trait values (Richardson and Rejmánek 2011, Sperry et al. 2021, Ni et al. 2021). Particularly on oceanic islands, island age (or soil age) - which typically has been used to address the macroevolutionary processes underlying native biodiversity patterns (Emerson 2002, Gillespie 2016, Whittaker et al. 2017) - may further strengthen the context dependency of biological invasions by differentially shaping the biodiversity patterns for native (Whittaker et al. 2008, Craven et al. 2019) and alien species (Leihy et al. 2018). Older islands with high native biodiversity may be more resistant to invasion (“Eltonian biotic resistance”; Fridley et al. 2007), yet there is growing evidence that more diverse native plant communities can also be more vulnerable to invasion (Stohlgren et al. 2003, Qian and Sandel 2022), possibly due to empty niche space, high habitat heterogeneity or greater resource availability. In the absence of disturbances, alien species are more apt to establish on older soils that usually have greater resource availability than on younger soils, which tend to be more nutrient poor (Aplet et al. 1998, Hughes and Denslow 2005, Zimmerman et al. 2008). This suggests that older soils may favor stronger biotic filtering, while younger soils may act as an environmental filter of biological invasions. Moreover, the context dependency of biological invasions could also be attributed to the phylogenetic composition of native island floras, which may affect the phylogenetic or functional similarity of alien species to native species, and therefore, biotic filtering (Bach et al. 2022). However, the extent to which soil age and anthropogenic and abiotic drivers underpin context-dependent biological invasions across oceanic islands remains uncertain.

Oceanic archipelagos, such as the Hawaiian islands, are frequently used as natural laboratories to disentangle ecological and evolutionary drivers of native biodiversity patterns (Emerson 2002, Gillespie 2016, Craven et al. 2019), yet also provide a compelling template for understanding biological invasions because of their recency and severity (Wohlwend et al. 2021, Barton et al. 2021). Here, we examine patterns of invasions of alien woody species, which represent approximately 60% of alien species on oceanic islands (Kueffer et al. 2010), and the relative importance of abiotic, biotic and anthropogenic drivers and their interactions. We first identify the abiotic and anthropogenic drivers of biological invasions (e.g., temperature, aridity, soil age, and the human influence index) in terms of alien species richness and abundance relative to native communities. We then evaluate the relative importance of phylogenetic distinctiveness and differences in traits of alien woody species to the native community along environmental and anthropogenic gradients for the establishment and dominance of alien woody species. Lastly, we test the sensitivity of the interactive effects of abiotic, biotic, and anthropogenic factors to the presence of the widespread and abundant native woody species, *Metrosideros polymorpha* (local name, Ohi’a), which is closely related to several alien and native woody species (Fig. S1; Craven et al. 2018). More specifically, the sensitivity analysis allows us to evaluate whether the relative importance of biotic filtering, *via* the phylogenetic and functional similarity of alien species to the native community, is due to a unique feature of native Hawaiian forests - the hyperdominance of *M. polymorpha* (Craven et al. 2018, Barton et al. 2021) - or a more general feature, such as their phylogenetic structure (Bach et al. 2022).

## Methods

### Plot data selection and preparation

To examine patterns of woody plant invasion of Hawaiian forests, we used a forest plot database that compiled data from multiple sources (Zimmerman et al. 2007, Ainsworth et al. 2011, Gillespie et al. 2013, Ostertag et al. 2014, US Forest Service 2016) to include species identity and size from across the Hawaiian archipelago (Craven et al. 2018). For our analyses, we used a total of 460 plots that range in area from 100 to 1,018 m^2^ (median plot area = 1,000 m^2^) with a minimum tree size threshold of 5 cm diameter at 1.3 m height. Plots were generally sampled to characterize community structure of Hawaiian forests, with one exception (Knight and Barton, unpublished; *n* = 35 plots) that focused specifically on forests including *M. polymorpha*. These plots are distributed across the six major islands in the archipelago: Hawai’i, Maui, Lana’i, Moloka’i, O’ahu, and Kaua’i (Fig. 1).

**Figure 1.**
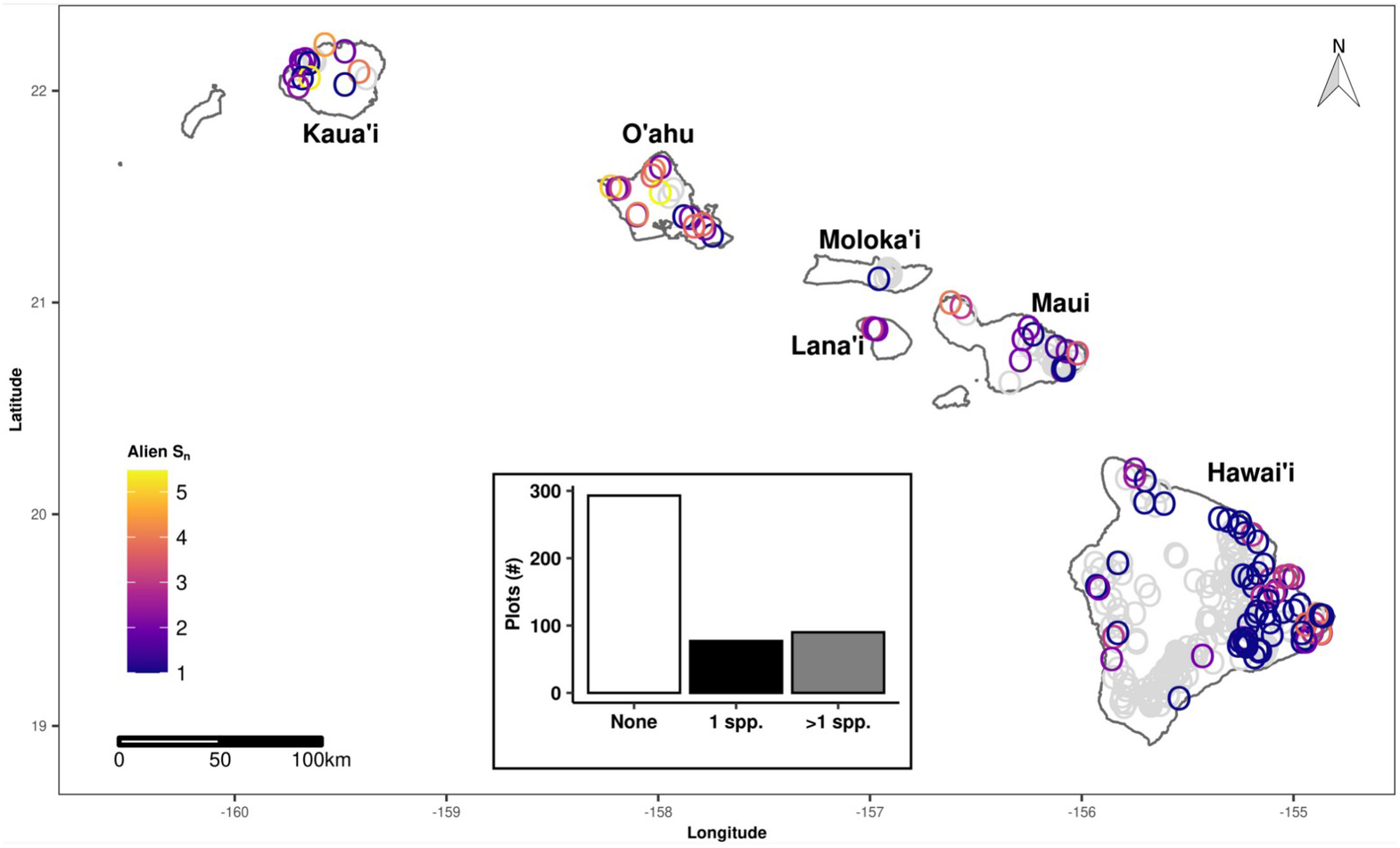
Map of alien species richness in forests across the Hawaiian archipelago n= 460 plots). Color corresponds to species richness of alien woody species rarefied to 100 stems; grey circles indicate plots that have not been invaded. Inset shows the number of plots invaded by zero, one, and more than one alien woody species; only individuals with a diameter greater than 5 cm at 1.3 m high were included.

### Species selection

Species names were standardized using The Plant List v 1.1 (www.theplantlist.org) with ‘Taxonstand’ (Cayuela et al. 2017) and alien status was obtained from the flora of the Hawaiian Islands (Wagner et al. 2020). While there are few non-angiosperm alien woody species in Hawai’i that have naturalized (*Pinus caribea* and *Cryptomeria japonica* (Wagner et al. 2020)), their branches in a phylogeny may be longer than those of angiosperms and may result in unusual phylogenetic measures (Qian et al. 2017). Similarly, monocot species likely have different ecological strategies than woody angiosperms (Spicer and Groover 2010, Pierce et al. 2013). We therefore excluded 14 non-angiosperm and monocot species from our dataset. We further excluded alien woody species that have not been reported as naturalised or whose populations are planted and have not spread widely, such as *Eucalyptus robusta* and *E. grandis* (Imada 2012, Wagner et al. 2020). Consistent with global patterns on oceanic islands (Delavaux et al. 2021), a large proportion of native and alien woody species are arbuscular mycorrhizal (90.9% and 82.9%, respectively; Soudzilovskaia et al. 2020) in our dataset (Fig. S1). In total, we included data for 129 woody species, of which 41 are alien and 88 are native (Table S1).

### Levels of biological invasion

We calculated commonly used metrics that are proxies for the establishment and dominance of alien species (Catford et al. 2012, Leung et al. 2012). At the community level, we calculated relative alien species richness as the proportion of alien woody species to all woody species, and relative alien species abundance as the proportion of alien woody stems to all woody stems as an indication of the extent of invasion. At the species level, we calculated presence/absence and relative abundances of each alien woody species. We classified absences as true absences if a species was reported as naturalized on an island by Wagner et al. (2020), but was not present in a plot. From here on, we refer to presence/absence as establishment and relative abundance (at the species level) as dominance.

### Phylogenetic distinctiveness of alien woody species

We used an updated version of the molecular phylogeny from Zanne et al. (Zanne et al. 2013, Qian and Jin 2016) to build phylogenies of all species (native and alien) and only alien species. We conservatively bound species to the backbone using dating information from congeners with the ‘congeneric.merge’ function in the ‘pez’ package (Pearse et al. 2015). In total, we placed all 129 woody species in the plot database on the phylogeny (Fig. S2).

Following Thuiller et al. (2010), we calculated phylogenetic distinctiveness of alien woody species relative to the community of woody native species using the mean abundance-weighted distance to the native community (MWDNC; phylogenetic distinctiveness from hereon), which is less sensitive to topological uncertainty in the phylogeny than other metrics, such as the phylogenetic distance to the nearest native species (DNNS; Park et al. 2020). Phylogenetic distinctiveness was calculated for each alien woody species in all plots, including those in which they were not present. Because phylogenetic distinctiveness may be confounded with species richness (Kembel 2009), phylogenetic distinctiveness is likely to be smaller in more diverse communities. To correct for this, we calculated a standardized effect size (SES) by shuffling the tips of the phylogeny 1,000 times:

SES = (observed - mean_null_) / standard deviation_null_.

Lower (more negative) SES values indicate that alien woody species are more closely related to the native plant community, while higher SES values indicate that alien woody species are less closely related to the native plant community.

### Functional traits

We selected three functional traits that are associated with a species’ competitive ability and that are associated with key axes of ecological variation (Díaz et al. 2016) to explore variation in establishment and dominance of alien woody species: wood density, adult plant size, and seed mass. Wood density is related to interspecific variation in growth and mortality rates (Chave et al. 2009), adult plant size is associated with competition for light (King et al. 2006) and seed dispersal distance (Thomson et al. 2011), and variation in seed mass drives the tolerance-fecundity trade-off (Muller-Landau 2010). Data for wood density were obtained from Chave *et al*. (2009) and from the BAAD (Falster et al. 2015), BIEN (Maitner 2020), FIA (Burrill et al. 2018), and TRY (Kattge et al. 2020) databases. For seed mass, we obtained data from Shiels (2011) and the BIEN (Maitner 2020) and TRY databases (Kattge et al. 2020). Because wood density and seed mass are strongly conserved phylogenetically (Chave et al. 2009, Igea et al. 2017), we calculated genus-level means when species-level data were not available instead of phylogenetically imputing these values (see next section; Tables S1 and S2). Adult plant size is often sample-size dependent, as smaller individuals are usually more abundant than larger ones (King et al. 2006). We therefore estimated adult plant size from plot data as the 95^th^ percentile of all diameters greater than the 10^th^ percentile of the maximum observed diameter (D95_0.1_) for species with 20 or more stems or as the 95^th^ percentile of diameter of all diameters for species with fewer than 20 stems (D95) following King et al. (2006). Both measures are independent of abundance, strongly correlated with maximum height, and are robust to outliers (King et al. 2006).

### Trait imputation

Because data were not available for all species in our data set (wood density: 5.4% unavailable; seed mass: <1% unavailable; adult plant size: 13.2% unavailable) and these gaps did not exceed 30% of our dataset (Penone et al. 2014), we used phylogenetic trait imputation to fill gaps in our data set. Including imputed trait data has also been shown to reduce bias compared to removing species with missing trait data (Penone et al. 2014). We used a random forest imputation algorithm in combination with phylogenetic information, which we optimized for each trait to minimize imputation error. For each trait, we assessed the imputation error for a range of phylogenetic eigenvectors (1 - 30) and used the imputed trait values for the number of phylogenetic eigenvectors that minimized imputation error. Prior to imputation, we log-transformed seed mass and adult plant size and back-transformed imputed values and imputation errors. We calculated imputation errors as the percent normalized root mean squared error, which were 1.95% for wood density, 13.6% for adult plant size, and 23.8% for seed mass. Our gap-filled trait data had similar distributions and mean values as the observed trait data (Fig. S3).

### Trait differences

To measure trait differences among native and alien species, we calculated differences between trait values of alien species and the abundance-weighted trait values of native species for each trait (Fig. S4; van der Sande et al. 2020). We standardized trait differences by calculating the log-response ratio of trait values of alien species to native species. Positive trait differences indicate that alien species had higher trait values than the native plant community and negative trait differences indicate the opposite.

### Abiotic conditions and anthropogenic impacts

To understand the role of abiotic conditions in shaping woody invasions, we extracted mean annual temperature (MAT), precipitation (MAP) and potential evapotranspiration (PET; calculated using the Penman-Monteith equation) for each plot from locally interpolated climate data (Giambelluca et al. 2013, 2014). We then calculated aridity, a widely used measure to assess the impacts of water availability on plant communities (e.g., Berdugo et al. 2020), as the ratio of MAP to PET (Zomer et al. 2008). We multiplied aridity by −1, such that low values of this index correspond to low aridity and high values to high aridity. We also extracted elevation (Jarvis et al. 2008) and soil substrate age for each plot (hereon referred to as soil age; Sherrod et al. 2007). On the Hawaiian archipelago, soil age is strongly related to the timing of volcanic activity over a volcanic hotspot, such that the western-most large islands (Kauai and O’ahu) are several millions of years older than the younger islands of Maui and Hawaii (Clague and Sherrod 2014).

Because the dispersal of alien woody species is facilitated by roads, and large populations of alien species can be maintained in areas where anthropogenic disturbance is high, we also included an index of human impacts in our analyses. Specifically, we extracted the human influence index (HII), a composite of human population density, land use, and infrastructure (HII; Wildlife Conservation Society - WCS and Center for International Earth Science Information Network - CIESIN - Columbia University 2005).

### Data analysis

Prior to model fitting, we tested for collinearity among explanatory variables using pairwise Pearson correlations. We selected variables that had a pairwise correlation below 0.7 (Fig. S5; Dormann et al. 2013). Based on this criteria, we retained mean annual temperature (MAT), aridity, soil age, and human influence index (HII) in our models examining relative alien species richness and abundance and MAT, aridity, soil age, HII, phylogenetic distinctiveness (in terms of mean weighted distance to native community (MWDNC)) and trait differences for wood density, adult plant size, and seed mass in our models examining variation in establishment and dominance. We opted to retain MAT instead of elevation in our models because it is a more direct measure of climatic conditions that is widely used to examine spatial patterns of alien plant invasions (e.g., Hulme 2017, Moser et al. 2018).

We examined the relative importance of aridity, MAT, HII, and soil age, in determining variation in relative alien species richness and abundance using hierarchical Bayesian models with Beta and lognormal distributions, respectively. Relative alien species richness and abundance were modeled as the proportion of alien woody species or individuals in a plot. In both models we used random intercept terms for island and plot size.

Because phylogenetically closely related species are likely to share similar trait values (Felsenstein 1985), not accounting for phylogenetic relationships may reduce estimation accuracy and increase type I error rates (Li and Ives 2017). We therefore fit phylogenetic hierarchical Bayesian models to examine variation in establishment and dominance of alien woody species as a function of abiotic, biogeographical, and anthropogenic factors (aridity, MAT, HII, soil age), phylogenetic distinctiveness, competitive trait differences, and two-way interactions of phylogenetic distinctiveness or trait differences with abiotic conditions. We fit the model for probability of occurrence (i.e. establishment) using a Bernoulli distribution and the model for the relative abundance of alien woody species using a lognormal distribution, which fits the data better than a Beta distribution. For both models, we used a crossed random effects structure with random intercept terms for island, species and plot, as not all alien species occurred in each plot or island. In the random intercept term for plot, we nested plot identity within a group factor of plot area to account for the increased probability of occurrence in larger plots. To account for phylogenetic correlations among species, we included two random intercept terms for species; one term models phylogenetic covariance and another term accounts for repeated measurements and other effects that may be independent of phylogenetic relationships among species. We first fit a full model with all two-way interactions and then simplified it by removing two-way interactions whose 95% credible intervals overlapped with zero.

We fit all models using weakly informative priors, four chains, and 2,500 burn-in samples per chain, after which 2,000 samples per chain were used to calculate posterior distributions of model parameters (total post-warmup samples= 8,000). Priors were flat for population-level effects. We increased the *‘adapt_delta’* parameter in the ‘brm’ function until there were no divergent transitions. Model convergence was evaluated visually with trace plots and by estimating *‘Rhat’* using the ‘rhat’ function, where values greater than 1.01 indicate that models have failed to converge (Tables S3 and S4). Model fit was assessed visually by comparing observed data to simulated data from the posterior predictive distribution (Fig. S6 and S7). Additionally, we estimated a Bayesian *r*^2^ for each model to represent an estimate of the proportion of variation explained for new data (Tables S3 and S4). Model fit for establishment (i.e., presence/absence) was also assessed by calculating ROC and AUC (Fig. S8). We evaluated spatial autocorrelation of model residuals using Moran’s I. All models showed minimal evidence of spatial autocorrelation (Figs. S9 and S10).

We tested all models for scale dependence by evaluating the precision and accuracy of model estimates per plot size (Spake et al. 2021); random effect estimates and their accuracy and precision did not vary systematically with plot size (Fig. S11 and S12). Additionally, we tested the sensitivity of our results to the presence of the widespread and abundant native woody species, *M. polymorpha*, by fitting the same set of models described above, but after removing *M. polymorpha* from our data set and phylogeny. We further tested whether mycorrhizal type contributed to understanding patterns of establishment or relative abundance of alien woody species by adding mycorrhizal type to our models as a categorical variable. We found that both establishment rates and relative abundance were similar across mycorrhizal types (Fig. S13), possibly because most of native and exotic species associate with arbuscular mycorrhiza (Fig. S1). We therefore decided that adding this variable to our models would not contribute to explaining patterns of establishment and dominance of alien woody species. All models were fitted using ‘brms’ (Bürkner 2017).

For all analyses, explanatory variables were visually inspected for normality; only soil age was natural-log transformed to ensure a linear relationship with response variables. All variables were standardized using a *z*-transformation to enable comparisons across models. Analyses were performed using R v. 3.6.1 (R Core Team 2020).

## Results

Woody invasions were widespread across Hawaiian forests on all six islands (Fig. 1). Approximately 36% of forest plots were invaded, 19.6% being invaded by multiple alien woody species and 16.7% being invaded by one alien woody species. Across all plots, the most frequent woody invaders were *Psidium cattleianum, Cecropia obtusifolia, Psidium guajava*, and *Melastoma malabathricum* (Fig. S2) and the most abundant woody invaders (present in at least five plots) were *Psidium cattleianum, Melastoma malabathricum, Schefflera actinophylla*, *Macaranga mappa*, and *Schinus terebinthifolia*.

### Abiotic and anthropogenic drivers of woody invasions

Relative species richness and abundance of alien woody species were moderately positively correlated (*r* = 0.39, P < 0.001). At the community level, invasion of forests by alien woody species varied with abiotic conditions (Fig. 2). Relative species richness and abundance of alien woody species increased with MAT, and relative abundance of alien woody species decreased with increasing aridity. Relative species richness and abundance of alien woody species also showed a tendency to increase with HII, as 80% credible intervals did not overlap with zero. We found no evidence for an effect of soil age on the richness or abundance of alien woody species. While the model for relative species richness of alien woody species explained 35% (95% credible interval: 26% - 44%) of its variation, the model for abundance of alien woody species only explained 18% (95% credible interval: 10%-26%) of its variation (Table S3).

**Figure 2.**
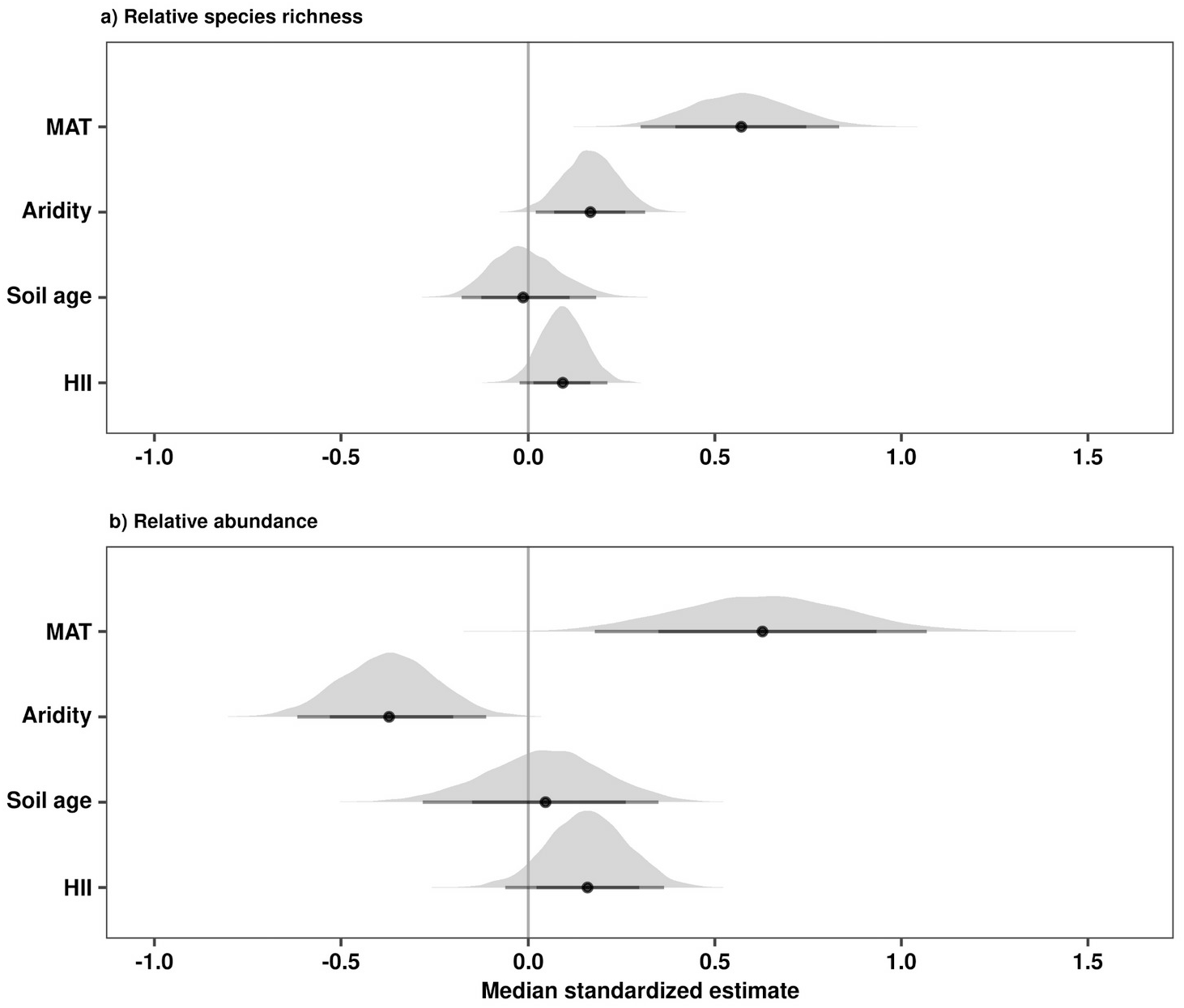
Coefficient estimates of main effects used to examine **a)** relative species richness and **b)** relative abundance of alien woody species across the Hawaiian archipelago. MAT is mean annual temperature, aridity is the ratio of mean annual potential evapotranspiration (PET) to mean annual precipitation, soil age is the estimated soil age, and HII is the human influence index. Medians, 80%, and 95% credible intervals of model coefficients were estimated using a phylogenetic hierarchical Bayesian with a beta distribution.

### Establishment of alien woody species

Our analysis showed that interactions among biotic and abiotic factors influenced the establishment of alien woody species across Hawaiian forests. Establishment of alien woody species increased with increasing temperature, soil age, and Δ adult plant size (Fig. 3) and the model explained 29% (95% credible interval: 25% - 32%; Tables S4 and S5) of its variation. Yet, all biotic factors exhibited differential responses along abiotic and anthropogenic gradients (Fig. 4). At higher temperatures, alien woody species more closely related to the native plant community were more likely to occur than those distantly related to the native plant community (Fig. 4a). Likewise, alien woody species with lighter seeds and smaller adult plant size than the native plant community were more likely to establish at higher temperatures (Fig. 4b and c). In areas with high aridity, alien woody species distantly related to the native plant community were more likely to occur than less distantly related ones, whose probability of establishment increased as aridity decreased (Fig. 4d). Alien woody species with heavier seeds and that are taller than the native plant community were more likely to occur where aridity was high, a pattern that switched sharply where aridity was low (Fig. 4e and f). Alien woody species with larger seeds and denser wood than the native plant community were more likely to occur on older soils than on younger soils (Fig. 4h and i). In areas with low human impacts, alien woody species with denser wood than the native plant community were more likely to establish than those with lighter wood than the native plant community (Fig. 4g). In contrast, alien woody species with lighter wood than the native plant community were more likely to establish in areas with higher human impacts (Fig. 4g).

**Figure 3.**
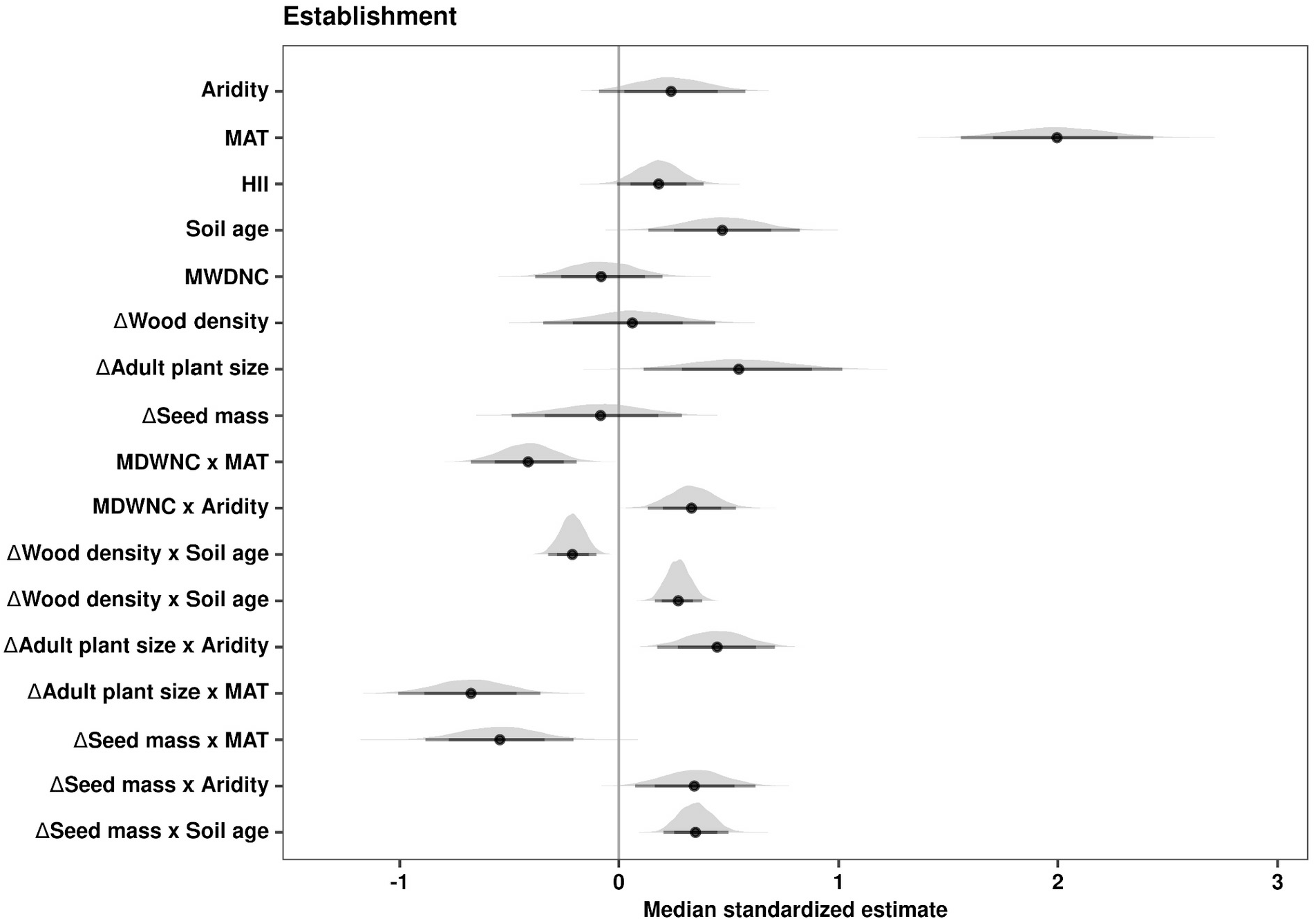
Coefficient estimates of main effects used to evaluate the establishment of alien woody species across the Hawaiian archipelago. Aridity is the ratio of mean annual potential evapotranspiration (PET) to mean annual precipitation, MAT is mean annual temperature, soil age is the estimated soil age, HII is the human influence index, MWDNC, i.e. phylogenetic distinctiveness, is the mean abundance-weighted phylogenetic distance to the native tree community, and ΔWD, Δ seed mass, and Δ adult plant size are the standardised differences in wood density, seed mass, and adult plant size, respectively, between alien woody species and the native community. Medians, 80%, and 95% credible intervals of model coefficients were estimated using a phylogenetic hierarchical Bayesian model with a Bernoulli distribution.

**Figure 4.**
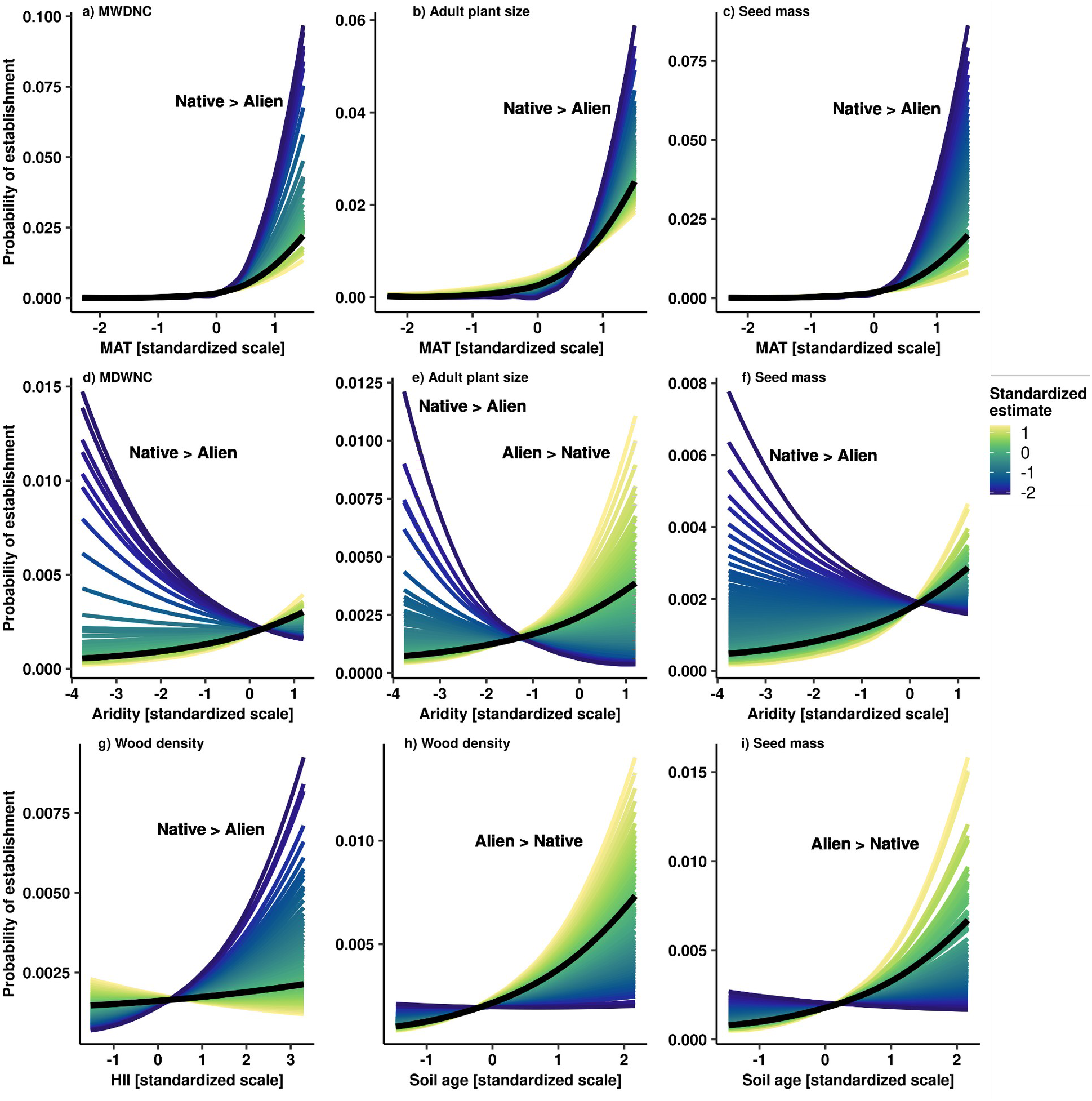
The estimated effects of phylogenetic distinctiveness (MDWNC) and trait differences among alien and native species in terms of seed mass, adult plant size, and wood density along gradients of (**a,b,c**) mean annual temperature (MAT), (**d, e, f**), aridity, (**g**) human influence index (HII), and (**h & i**) soil age on the probability of establishment of alien woody species across Hawaiian forests. MWDNC is the mean abundance-weighted phylogenetic distance to the native tree community; positive values indicate that alien species are phylogenetically distinct from the native plant community and negative values indicate that alien species are phylogenetically similar to the native plant community. Seed mass, adult plant size, and wood density are standardized trait differences between alien woody species and the native community; positive values indicate that trait values of alien species are larger than those of the native community and negative values indicate that trait values of the alien woody species are smaller than those of the native community. Black and colored lines are marginal predictions evaluated at the 50th quantile and from the 5th to the 95th quantile of the moderator, respectively, and estimated with a phylogenetic hierarchical Bayesian model with a Bernoulli distribution. All x axes are scaled to unit variance.

### Dominance of alien woody species

Abiotic and biotic factors and their interactions did not have a marked influence on dominance patterns of alien woody species (Fig. 5). Only phylogenetic distinctiveness had a moderate, negative effect on the relative abundance of alien woody species (80% credible intervals did not overlap zero), and no interactions between abiotic and biotic factors were included in the final model, which explained 47% of variation in dominance (95% credible interval: 35% - 57%; Tables S4 and S6), as the 95% credible intervals of all two-way interactions overlapped with zero.

**Figure 5.**
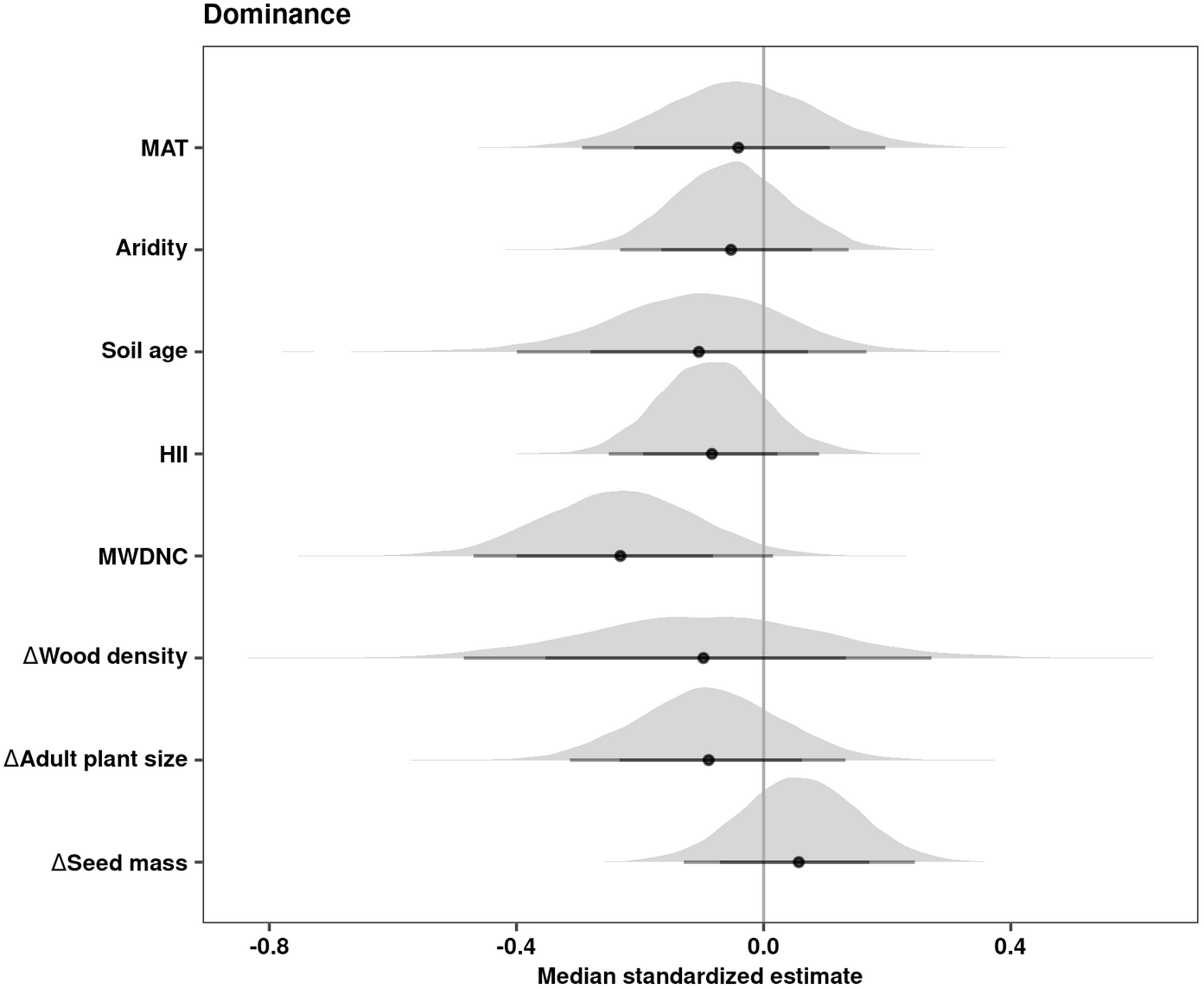
Coefficient estimates of main effects used to examine dominance of alien woody species across Hawaiian forests. Aridity is the ratio of mean annual potential evapotranspiration (PET) to mean annual precipitation, MAT is mean annual temperature, soil age is the estimated soil age, HII is the human influence index, MWDNC, i.e. phylogenetic distinctiveness, is the mean abundance-weighted phylogenetic distance to the native tree community, and ΔWD, Δ seed mass, and Δ adult plant size are the standardised differences in wood density, seed mass, and adult plant size, respectively, between alien woody species and the native community. Medians and 80% and 95% credible intervals of model coefficients were estimated using a phylogenetic hierarchical Bayesian model with a lognormal distribution.

### Sensitivity of context-dependent drivers of establishment and dominance

Further analyses provided added insight to the role played by *M. polymorpha* in mediating the context dependency of drivers affecting the establishment and dominance of alien woody species. When we removed *M. polymorpha* from the dataset, the influence of many interactive effects identified in the main analysis on the establishment of alien woody species was reduced considerably (Fig. S14). Indeed, only the interactions between soil age and Δ seed mass, MAT and Δ seed mass, and HII and Δ wood density retained their influence on the probability of establishment of alien woody species. Independent of the presence of *M. polymorpha*, temperature, soil age, and Δ adult plant size exhibited strong, positive effects on the probability of establishment of alien woody species, while Δ seed mass had strong, negative effects on the probability of establishment of alien woody species.

In contrast, the removal of *M. polymorpha* from the data set resulted in minimal changes to the impacts of abiotic and biotic drivers on the dominance of alien woody species (Fig. S15). The exception to this was Δ seed mass, which had a positive effect on dominance. As found for the analysis of the full data set, no interactive effects were detected and phylogenetic distinctiveness had a marginally negative effect on the dominance of alien woody species.

## Discussion

Our results highlight the context dependence of biological invasions (e.g., Catford et al. 2022) in forests across the Hawaiian archipelago. Abiotic factors influence the richness and abundance of alien species in communities in the expected direction, with more woody aliens in hotter and wetter locations.

Interactions among biotic, abiotic, and anthropogenic factors jointly determine where alien woody species establish, but not where they become dominant. We found that alien woody species establish more readily in areas with high aridity and in older soils, which they most likely colonize successfully by exploiting niche space not occupied by native woody species. In areas with harsh abiotic conditions, such as those with younger soils, environmental filtering was usually - but not always - the likely dominant ecological mechanism operating on the establishment of alien woody species. Our results further reveal how the impacts of soil age, a biogeographical factor often used to examine macroevolutionary processes that generate native biodiversity across and within oceanic islands (Emerson 2002, Whittaker et al. 2008, Gillespie 2016), may extend to shaping local coexistence of alien species via its effects on native diversity (“Eltonian biotic resistance”). Together, our results suggest that the ecological mechanisms underlying both the pre-adaptation hypothesis (environmental filtering; Cadotte et al. 2018, Bennett 2019) and Darwin’s naturalisation conundrum (biotic filtering; Cadotte et al. 2018, Bennett 2019) are interactively shaping the assembly of alien woody communities across the Hawaiian archipelago.

### Abiotic drivers of multi-species woody invasions

We found that local communities are generally more invaded in forests with higher temperatures and low aridity. These findings indicate that biological invasions are more severe in lowland, moist tropical forests across the Hawaiian archipelago (Ibanez et al. 2019), which have been heavily impacted by agriculture and land-use change (Jacobi et al. 2017), as temperature is strongly negatively correlated with elevation (*r* = - 0.99). While our results show certain similarities with macroecological patterns of alien species richness and composition on oceanic islands in terms of abiotic factors, we detected weaker human impacts on the severity of biological invasions at local scales than have been reported at larger spatial scales (Moser et al. 2018, Wohlwend et al. 2021). Yet, the low explanatory power of our models for relative alien species richness and abundance highlights the need to include additional drivers, such as biotic interactions, historical legacies, and propagule pressure, to enhance the robustness of predictions of the severity of biological invasions at local spatial scales. Including small-statured shrubs which were not counted in the tree plots that we used (< 5 cm at 1.3 m high), such as the highly invasive shrub, *Clidemia hirta* (e.g., DeWalt et al. 2004), may further increase the explanatory power of these models by including a fuller picture of invasions by alien woody species in Hawaiian forests. Moreover, the low explanatory power of these models could also highlight the importance of stochastic processes, such as long-distance dispersal, in determining patterns of relative alien species richness and abundance.

The weak, positive correlation between the two metrics of invasion impacts suggests that while in general there are more alien species when there are more alien stems, there is also a range of invaded forests across Hawai’i, including those that are heavily invaded by a small number of dominant alien woody species such as *P. cattleianum* and those dominated by native woody species, but contain several alien woody species. This finding also suggests that invasion by one species may not facilitate invasion by other alien species within the same trophic group (e.g., Zarnetske et al. 2013), possibly due to niche preemption or niche modification (Fukami 2015). As such priority effects are often contingent upon abiotic factors (Grainger et al. 2019), it is possible that high temperatures facilitate coexistence among alien woody species, while low aridity may strengthen priority effects. Further studies would be needed to determine whether and how priority effects of dominant alien species such as *P. cattleianum* are influenced by their demographic traits (Fukami et al. 2016) to better understand the mechanisms that underpin multi-species biological invasions.

### Context dependency of alien species establishment and dominance

Our analysis suggests that the establishment of alien woody species was context dependent (sensu Catford et al. 2022), meaning that the impacts of phylogenetic and trait differences between alien and native species on establishment were not consistent across Hawaiian forests (Tecco et al. 2010, Gross et al. 2013, González-Moreno et al. 2014, Sapsford et al. 2020). The strength of biotic filtering is expected to shift along environmental gradients (e.g., Cornwell and Ackerly 2009, Kraft and Ackerly 2010), with the expectation that biotic filtering should be stronger in more favorable environments compared to harsher environments (Bertness and Callaway 1994). In contrast, we found that environmental filtering strengthened with increasing temperature (for phylogenetic distinctiveness, Δ seed mass, and Δ adult plant size), meaning that alien woody species that are more similar to native woody species are more likely to establish where environmental filters are weaker. In line with expectations, we found that environmental filtering strengthened with increasing human impacts (for Δ wood density) and decreasing soil age (for Δ seed mass and Δ wood density); alien woody species that are more similar to native woody species were more likely to establish in areas with stronger human impacts and/or with younger soils. Yet, we also found evidence that biotic filtering could be stronger in harsh environments, particularly where aridity was high (for phylogenetic distinctiveness, Δ adult plant size, and Δ seed mass). This is consistent with results reported by Henn et al. (2019). This finding suggests that native woody communities do not fully occupy niche space in harsh environments, thereby facilitating the establishment of phylogenetically and functionally distinct alien woody species. It could also suggest that alien woody species may exhibit greater tolerance to water- or nutrient-limited conditions than phylogenetically similar native woody species, which could be facilitated by greater intraspecific trait variation (Westerband et al. 2021). Together, our results reveal that alien woody species are likely to be able to colonize native forests by competing less strongly with native woody species (Levin et al. 2020), either because conditions are too harsh (i.e. where environmental filtering is strong) or empty niche space is available in places where biotic filtering is strong.

Our analysis shows that soil age is an important underlying factor driving context dependent biological invasions across Hawaiian forests. One explanation for the positive effects of soil age on the probability of establishment of alien woody species observed in this study is that woody communities with high native plant diversity on older soils are less resistant to invasion (“the rich get richer”; Stohlgren et al. 2003, Fridley et al. 2007). This suggests that particular soil properties associated with older soils (not measured in this study) may facilitate greater species richness of both native and alien woody communities. In line with this idea, we found evidence that the establishment of alien woody species increases in areas with older soils by occupying different regeneration niches or acquiring soil nutrients differently than native woody species (Fig. 4h and i), strategies that may covary with either wood density or seed mass (but see Carmona et al. 2021). Our results are also consistent with the idea that soil age directly affects biological invasions by filtering alien woody species on younger soils that have low soil nutrient content (Fig. 4h and i; Aplet et al. 1998). Moreover, once accounting for soil age, native and alien species richness exhibited a significantly negative correlation (Pearson’s partial correlation = - 0.11, *P* =0.01), indicating that soil properties associated with soil age may lower the resistance of native woody communities to biological invasions (Fridley et al. 2007). Therefore, soil age - *via* its effects on soil properties - likely plays a crucial role in determining where biological invasions occur in oceanic islands.

We found little evidence that abiotic or biotic filtering consistently influenced the dominance of alien woody species across Hawaiian forests. Our results indicate that competitive outcomes between alien and native woody species are largely independent of environmental conditions or functional or phylogenetic differences. Thus, spatial variation in dominance of the alien woody community could be driven principally by dispersal, particularly if propagule pressure is high (Lockwood et al. 2005, Richardson and Pyšek 2012, Cassey et al. 2018). We detected a moderate negative effect of phylogenetic distinctiveness on dominance (80% confidence intervals do not overlap with zero), which is consistent with the idea that environmental filtering is the dominant process structuring invaded forest ecosystems (Lemoine et al. 2015). Nevertheless, our ability to detect the influence of abiotic and biotic filtering on dominance may have been limited by the spatial grain of our analysis (median plot area = 1,000 m^2^; range of plot area: 100 to 1,018 m^2^), which may be too large to capture the effects of neighborhood competition or crowding (e.g., Uriarte et al. 2010). Further, we used phylogenetic distinctiveness and trait differences as proxies for competitive ability, which is rarely quantified directly (but see Levin et al. 2020). These may explain further variation in dominance among alien woody species, as could the inclusion of additional functional traits, such as demographic and root traits, that provide insights to other dimensions of plant form and function (Rüger et al. 2018, Carmona et al. 2021).

### Presence of M. polymorpha

We found that the patterns of alien establishment critically depend on the presence of *M. polymorpha*. When we excluded *M. polymorpha* from the analysis, the factors that shape the establishment of alien woody species were markedly less context dependent; only three interactions (soil age and Δ seed mass, MAT and Δ adult plant size, and HII and Δ wood density) were retained. This result suggests that where temperature and human impacts are high, alien woody species that are phylogenetically and functionally similar to native species are more likely to establish; this is likely driven by the pervasiveness of *P. cattleianum* across Hawaiian forests and its functional similarity to *M. polymorpha* (Fig. S16). It also suggests that independent of the presence of *M. polymorpha*, soil age mediates the strength of biotic filtering. Thus, the apparent context dependency of biological invasions appears to have been driven at least in part by the broad environmental gradients across which *M. polymorpha* occurs, whose light seeds allow it to disperse widely and high wood density confers greater drought tolerance (Cordell et al. 1998), as well as the functional similarities between alien woody species and *M. polymorpha*.

Our sensitivity analysis indicates that unoccupied niche space persists even in the absence of *M. polymorpha*, as evidenced by retained interactions (soil age *x* Δ seed mass, MAT *x* and Δ adult plant size, and HII *x* Δ wood density). These results suggest that being taller, having lighter seeds or denser wood than the native community in areas with older soils or lower human impacts may confer a competitive advantage to alien woody species invading forests, regardless of whether they are dominated by *M. polymorpha*. Consequently, the growing impacts of the fungal pathogen that causes Rapid O’hia Death on *M. polymorpha* could shift alien woody invasions in Hawaiian forests (Gómez-Aparicio et al. 2012, Anderegg et al. 2015), potentially favoring alien woody species with lighter seeds and taller stature than the native woody community, such as *Aleurites moluccanus*, *Melia azedarach*, and *Syzygium cumini*.

### Conclusions

Our results highlight that alien woody species likely establish in island ecosystems by occupying empty niche space and/or habitats that are too harsh to compete directly with native woody species. While these alien species are clearly changing the composition and dynamics of Hawaiian forests (Craven et al. 2019, Barton et al. 2021), our results show that biological invasions are shaped in part by the same environmental and biogeographical factors, such as climate and soil age, that generate and maintain patterns of native biodiversity. Our sensitivity analysis provides evidence that the context dependency of alien woody invasions across Hawai’i arises due to ecological interactions between abiotic, anthropogenic, and biotic factors (Catford et al. 2022), particularly along gradients in soil age, human impacts, and temperature. While this context dependency makes our results difficult to generalize to other archipelagos, our results emphasize that future efforts should account for how soil age mediates biotic filtering. This strong context dependence of multi-species invasions also highlights the complexity of developing management strategies to mitigate the biodiversity and ecosystem impacts of biological invasions.

## Supporting information

Supplementary materiales

## Acknowledgments

DC, TMK and JMC acknowledge funding by German Centre for Integrative Biodiversity Research (iDiv) Halle-Jena-Leipzig, funded by the German Research Foundation (FZT 118). DC also received funding from the Agencia Nacional de Investigación y Desarrollo (Chile; FONDECYT Regular No. 1201347). TMK also acknowledges support from the Alexander von Humboldt Foundation, and the National Geographic Society. We thank Valentin Stephan for helping to prepare the environmental data, Nathaly Guerrero-Ramírez, Patrick Weigelt, Gunnar Keppel, Thomas Ibanez, and Holger Kreft for their input to conceptual development, and to data providers for collecting the field data. We also thank Rebecca Spake for her feedback on data analysis. The authors certify that they do not have any conflict of interest.

## Data availability statement

The code and data that support the findings of this study are available in Github: https://github.com/dylancraven/Hawaii_BiologicalInvasions. These data were derived from the following resources available in Dryad: https://doi.org/10.5061/dryad.1kk02qr.

