## Supplementary materiales for "Interacting abiotic and biotic drivers shape woody invasions across Hawaiian forests"

This file includes:

Figures S1 to S16
Table S1 to S6

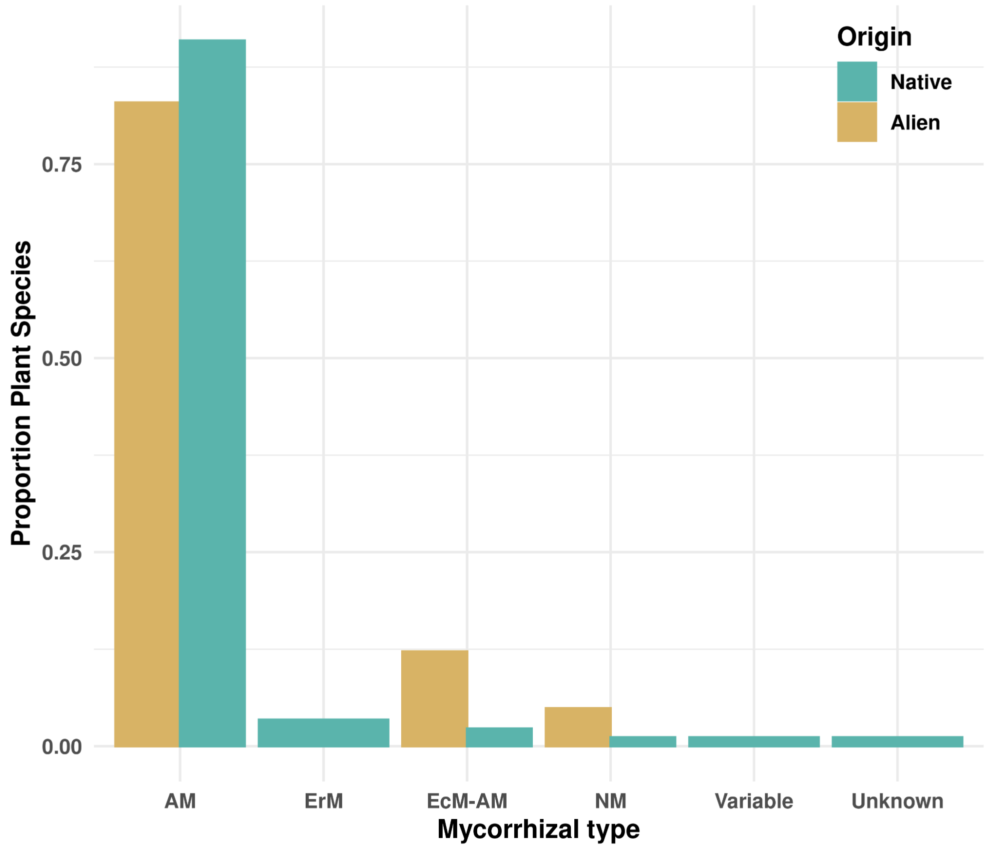


**Figure S1**. Proportion of native and alien species by mycorrhizal type across Hawaiian forests. Mycorrhizal types were assigned by plant genera; AM is arbuscular mycorrhiza, ErM is ericoid mycorrhiza, EcM-AM is ectomycorrhiza and arbuscular mycorrhiza, NM is no mycorrhiza, variable indicates that mycorrhiza type varies within a genus, and Unknown indicates that the mycorrhizal type is not known for a given plant genus.

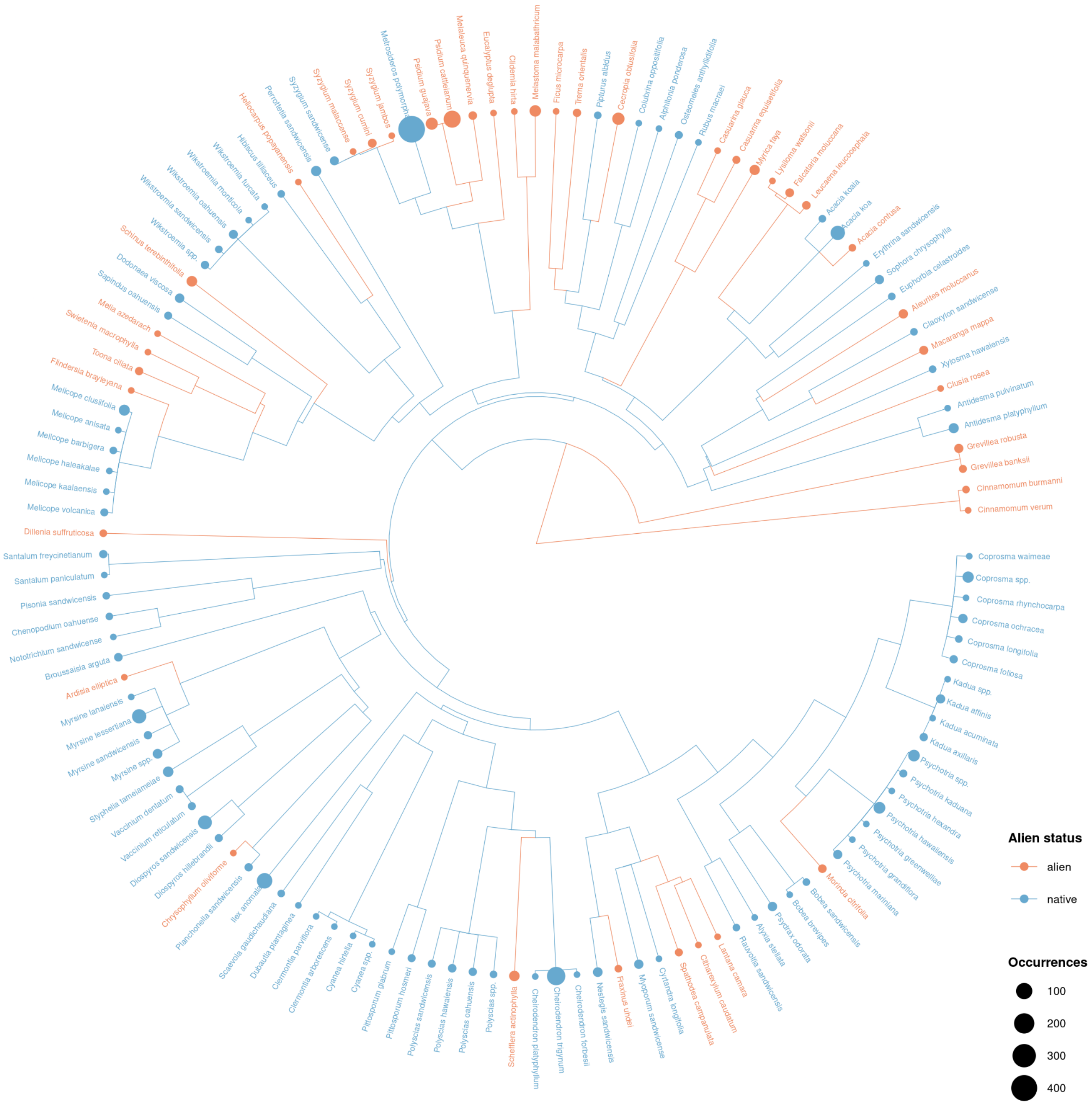


**Figure S2**. Phylogenetic structure of native and alien woody species in Hawaiian forests. Orange dots represent alien species (*n* = 41 species) and blue dots indicate native species (*n*= 88 species). Dots are scaled to the number of plots in which each species occurs.

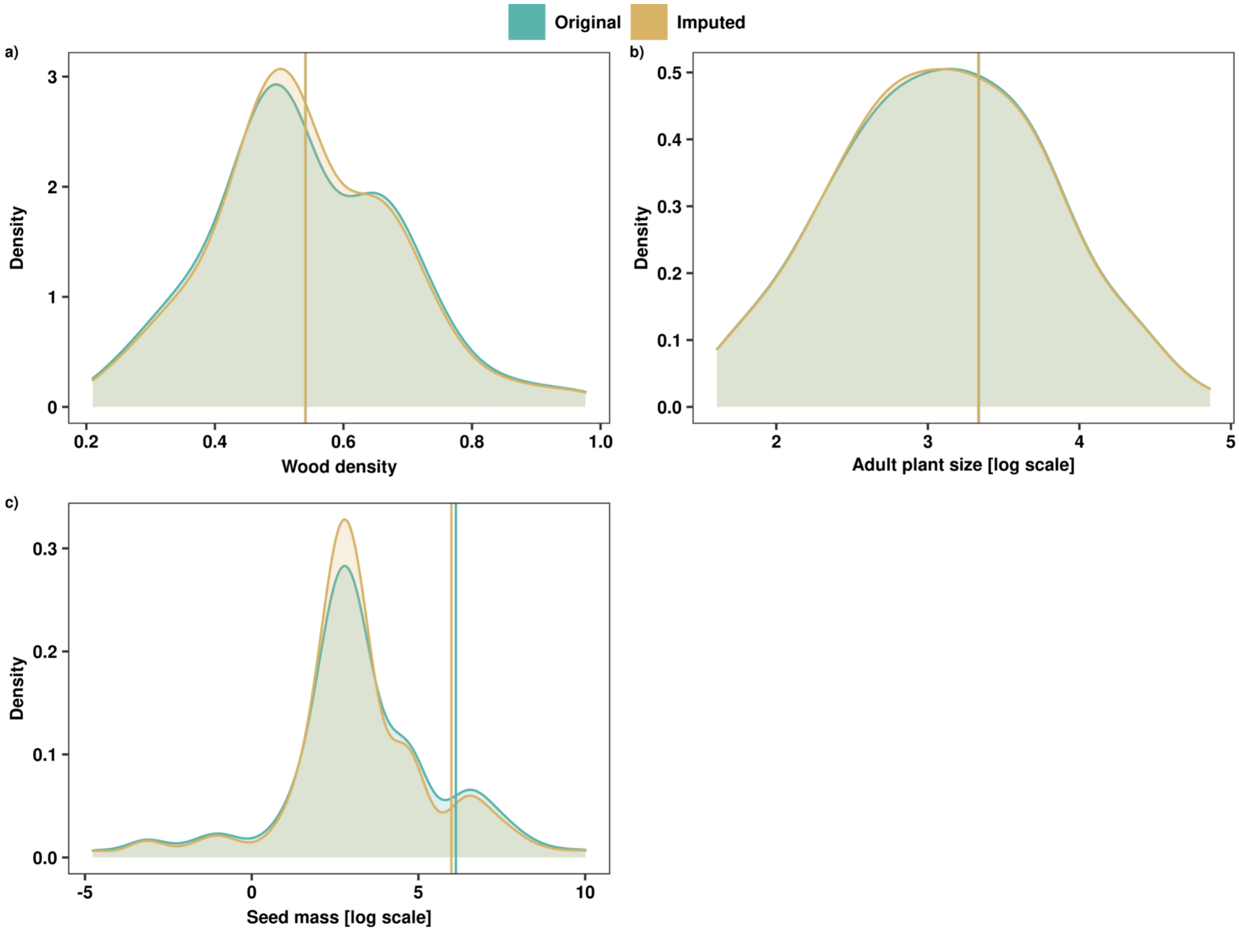


**Figure S3.** Comparison of distributions of original and imputed traits for wood density, adult plant size, and seed mass of 129 native and alien woody species. Trait values were imputed using phylogenetic information and a random forest algorithm.

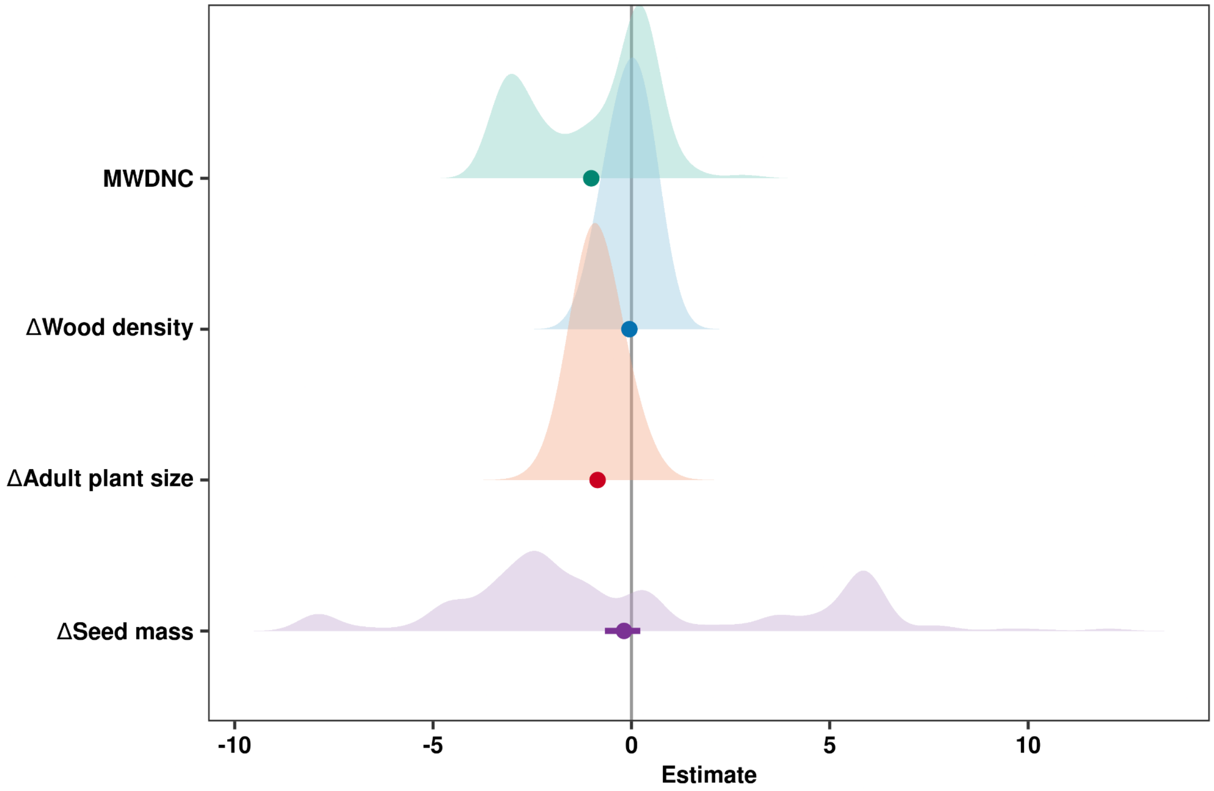


**Figure S4**. Summary of phylogenetic distinctiveness and competitive trait differences between alien and native woody species in Hawaiian forests. MWDNC, i.e. phylogenetic distinctiveness, is the standardized effect size of the mean weighted phylogenetic distance to the native tree community, and ΔWood density, Δ Seed mass, and Δ Adult plant size are the standardised differences in wood density, seed mass, and adult plant size, respectively, between alien woody species and the native tree community species. Points are means and lines are 95% bootstrapped confidence intervals. More negative MWDNC values indicate that alien species are more phylogenetically similar to native species, while more positive MWDNC values indicate that alien species are more phylogenetically dissimilar to native species. Positive trait differences indicate that alien species had higher trait values than native species and negative trait differences indicate the opposite.

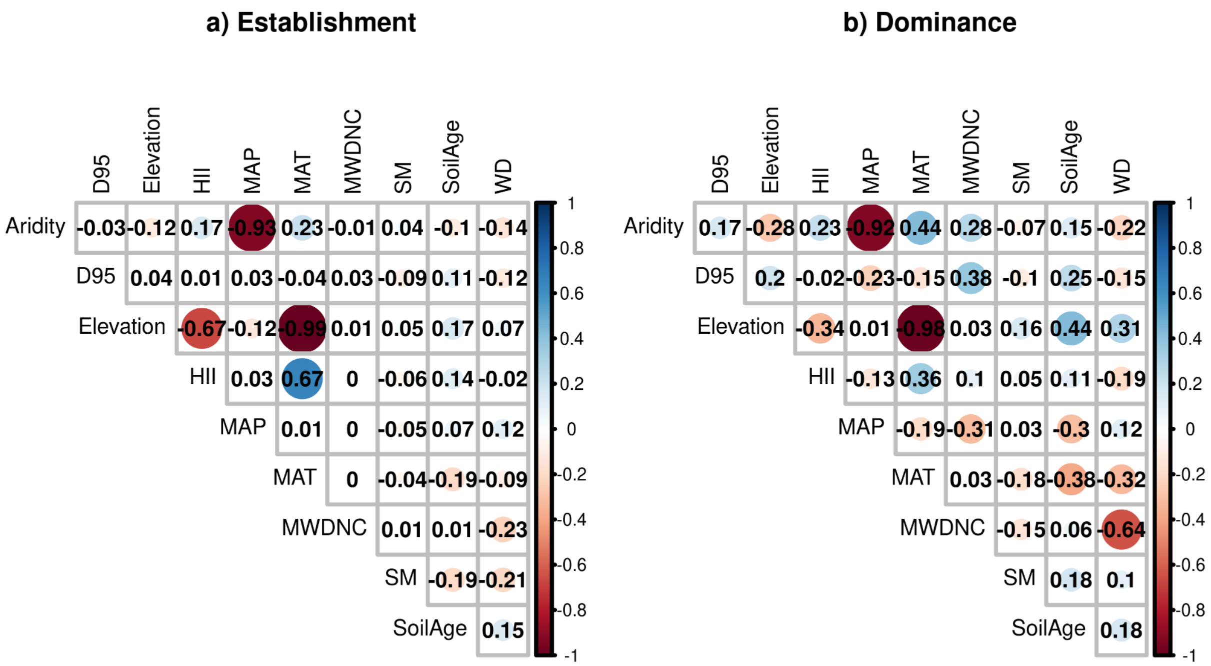


**Figure S5.** Pairwise correlations of potential explanatory variables to model **a**) establishment (i.e. presence/absence) and **b**) dominance (i.e. relative abundance) of alien woody species across Hawaiian forests. Aridity is the ratio of MAP to potential evapotranspiration (PET), D95 is the standardised difference in adult plant size between alien woody species and the native tree community, HII is the human influence index, MAP is mean annual precipitation, MAT is mean annual temperature, MWDNC, i.e. phylogenetic distinctiveness, is mean abundance-weighted distance to the native community, SM is the standardised difference in seed mass between alien woody species and the native tree community, SoilAge is the estimated soil age, and WD is the standardised difference in wood density between alien woody species and the native tree community. Soil age was natural-log transformed to meet normality assumptions.

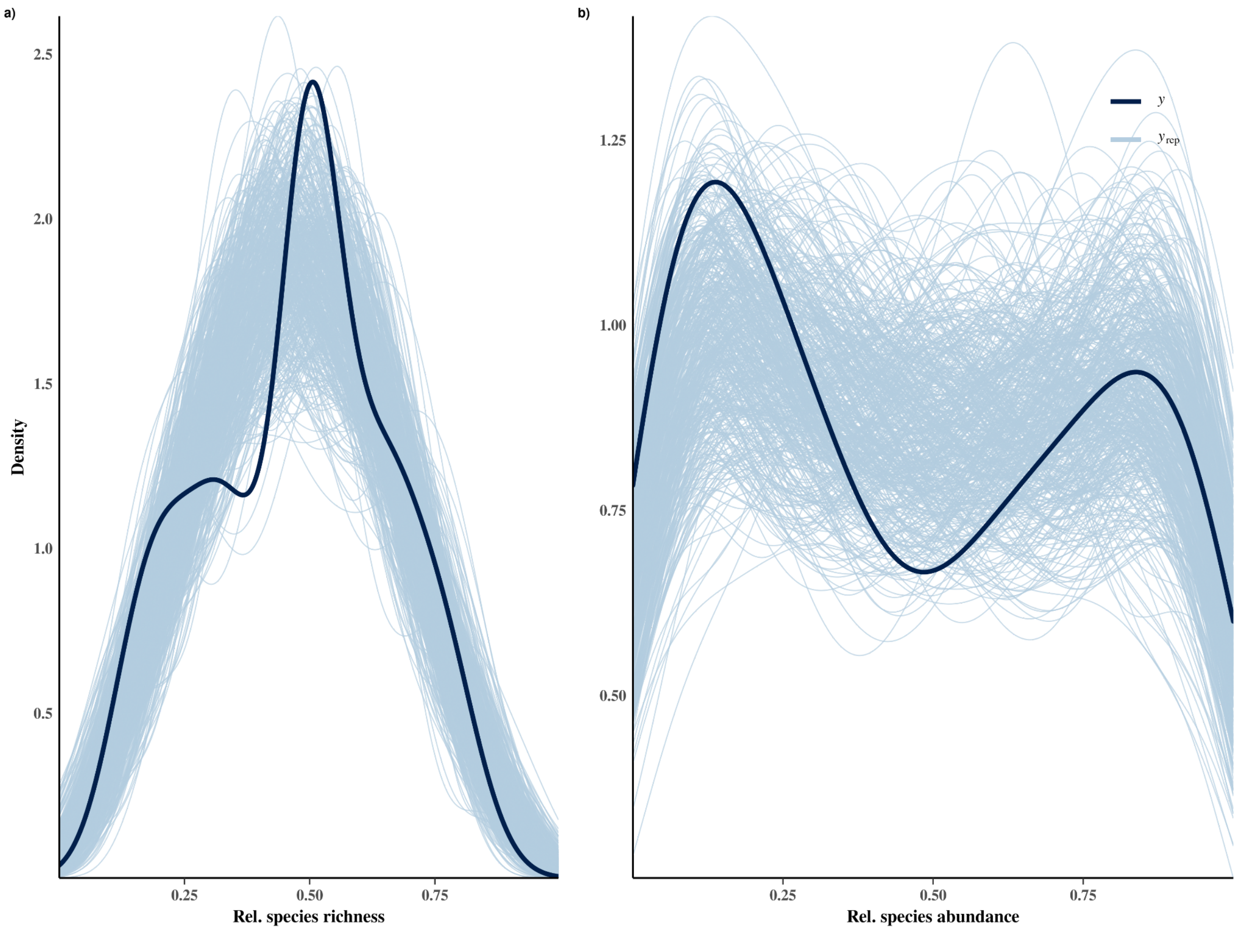


**Figure S6**. Posterior predictive distributions for hierarchical Bayesian models estimating **a**) relative species richness and **b**) relative abundance of alien woody species across Hawaiian forests. Dark blue lines represent the observed data, while the light blue lines represent simulations of the response variable (n = 500 posterior samples). Models for a and b were fit with a Beta distribution.

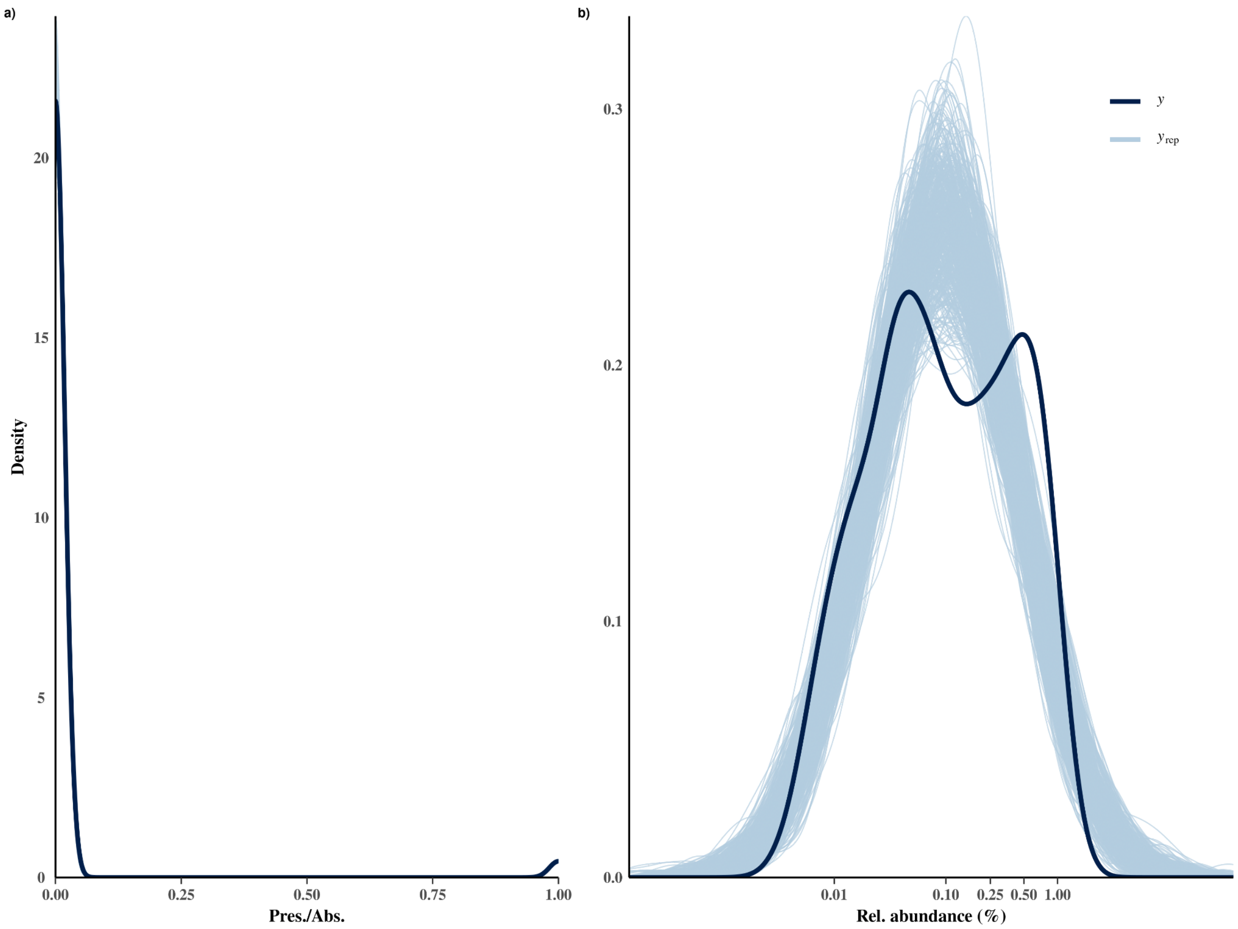


**Figure S7.** Posterior predictive distributions for phylogenetic hierarchical Bayesian models estimating **a)** establishment (presence/absence) and **b)** dominance (relative abundance) of alien woody species across Hawaiian forests. Dark blue lines represent the observed data, while the light blue lines represent simulations of the response variable (*n* = 500 posterior samples). The x-axis for **b**) is on the natural-log scale, as the corresponding model was fit using a lognormal distribution.

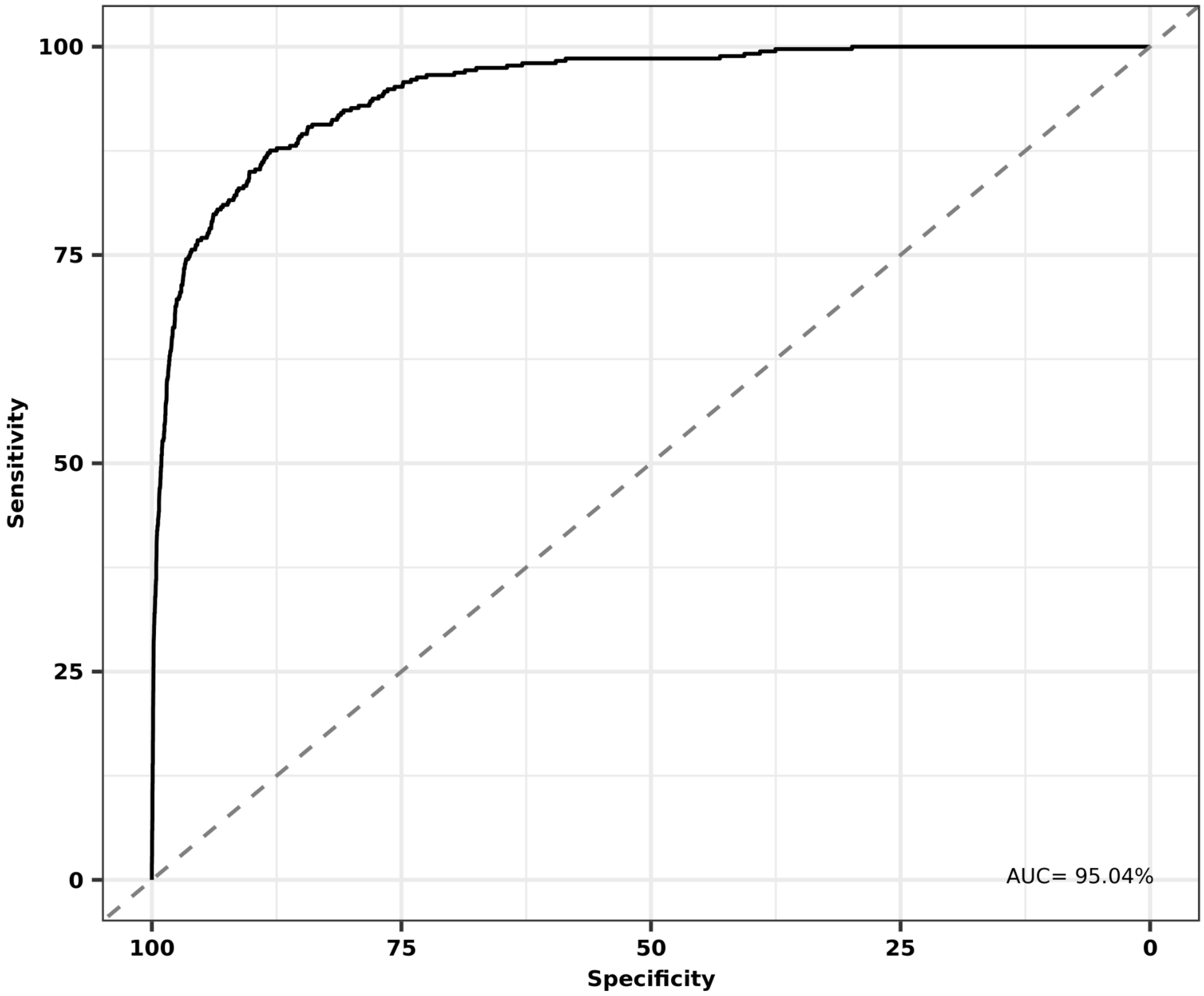


**Figure S8**. Receiver operating characteristic (ROC) curve for a phylogenetic hierarchical Bayesian logistic model examining the drivers of establishment of alien woody species across Hawaiian forests. Area under the curve (AUC) is also provided; ROC curves and AUC indicate the ability of binomial hierarchical models to make correct predictions.

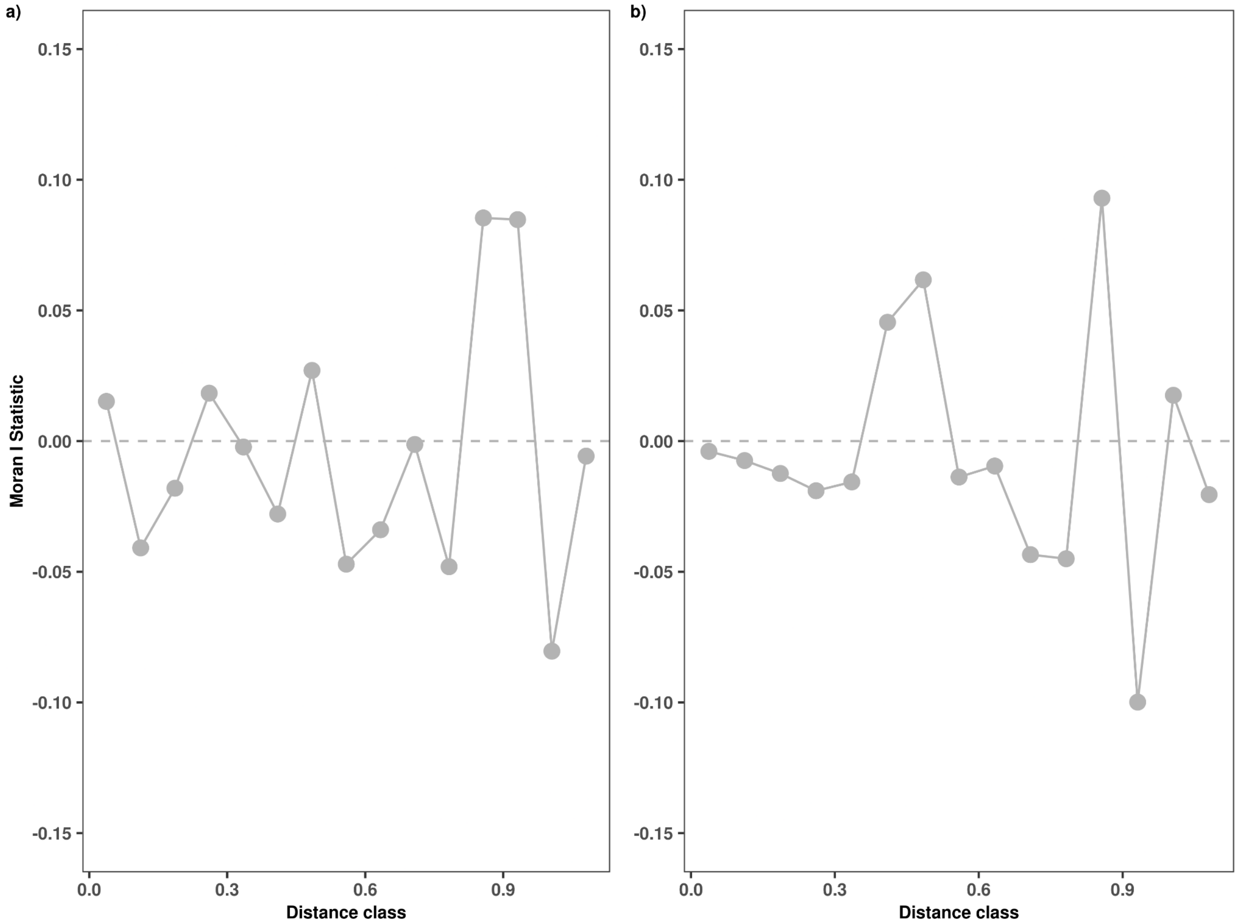


**Figure S9**. Spatial autocorrelation of model residuals for hierarchical Bayesian models that examine **a**) relative species richness and **b**) relative abundance of alien woody species across Hawaiian forests. Moran's I was calculated for 15 distance classes.

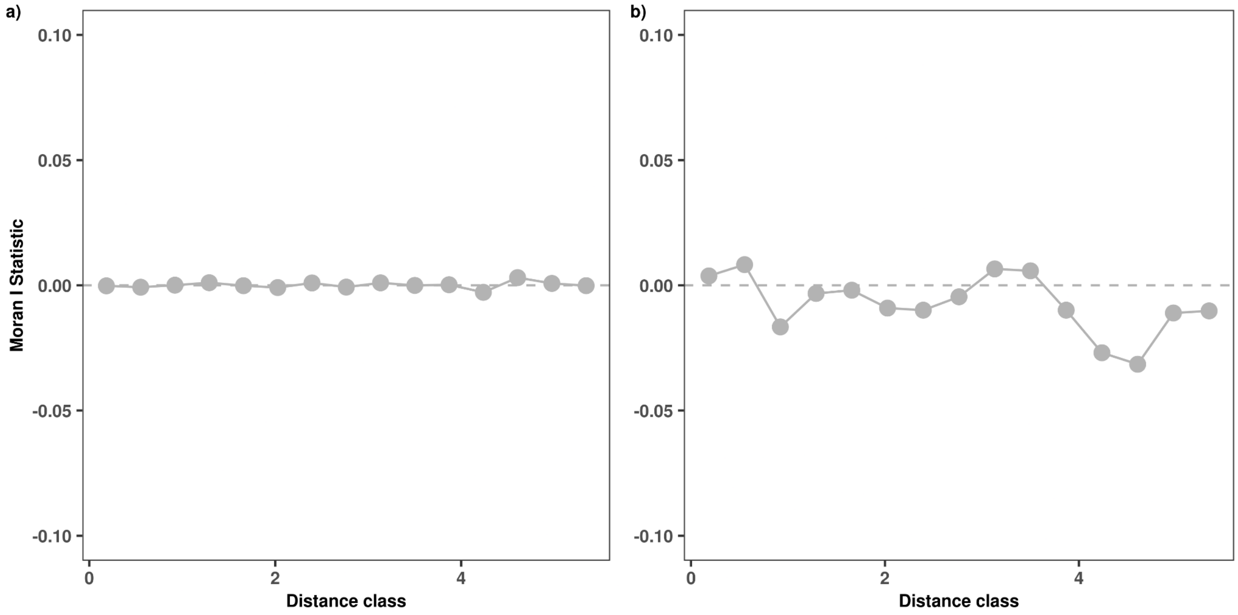


**Figure S10**. Spatial autocorrelation of model residuals for phylogenetic hierarchical Bayesian models that examine **a)** the establishment (presence/absence) and **b)** dominance (relative abundance) of alien woody species across Hawaiian forests. Moran's I was calculated for 15 distance classes.

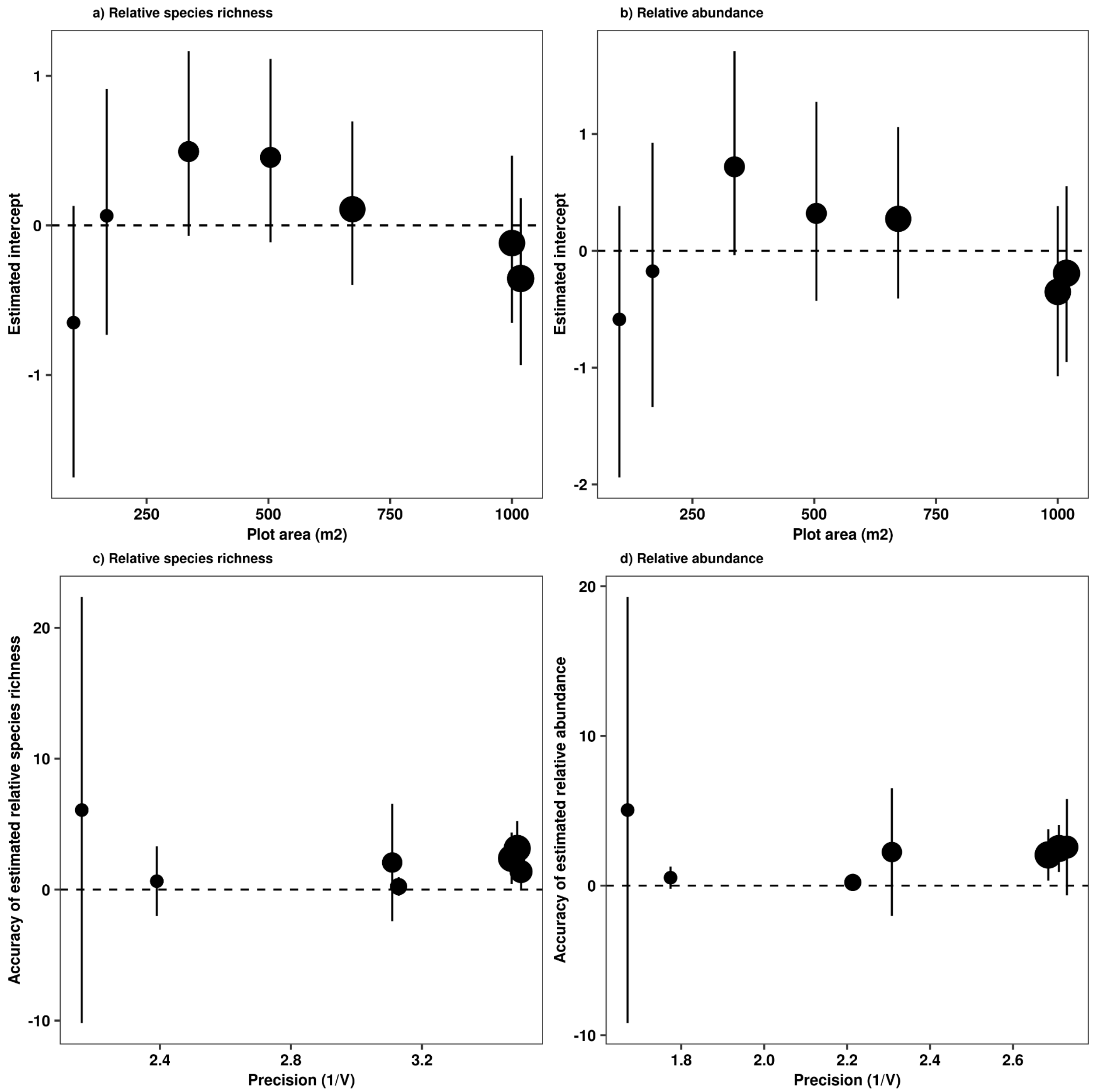


**Figure S11.** Relationships of estimated intercepts and plot area (**a** and **b**) and accuracy to precision across plot areas for relative abundance and relative species richness of alien woody species at the community level. Estimated intercepts are the random effects for plot area for each model, accuracy is the standardised difference of the estimated from the true value of the response variable (relative species richnessd or relative abundance), and precision is variation around the estimated response variable. Point size is scaled by the number of replicates for each plot size.

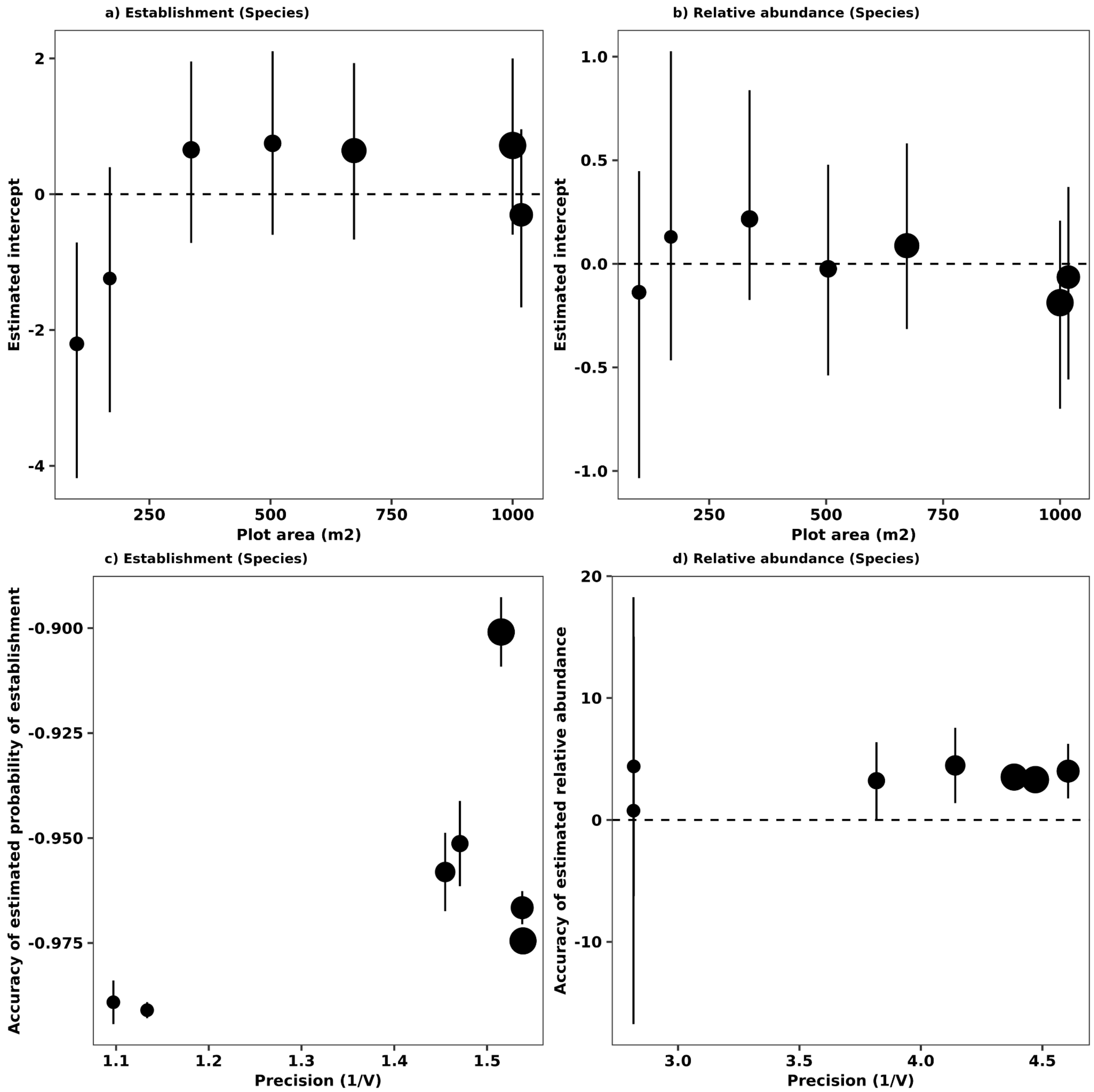


**Figure S12.** Relationships of estimated intercepts and plot area (**a** and **b**) and accuracy to precision across plot areas for establishment (presence/absence) and relative abundance of alien woody species at the species level. Estimated intercepts are the random effects for plot area for each model, accuracy is the standardised difference of the estimated from the true value of the response variable (establishment or relative abundance), and precision is variation around the estimated response variable. Because a Bernoulli distribution was fitted for the model estimating establishment, a 0.5 was used as the threshold to estimate accuracy. Point size is scaled by the number of replicates for each plot size.

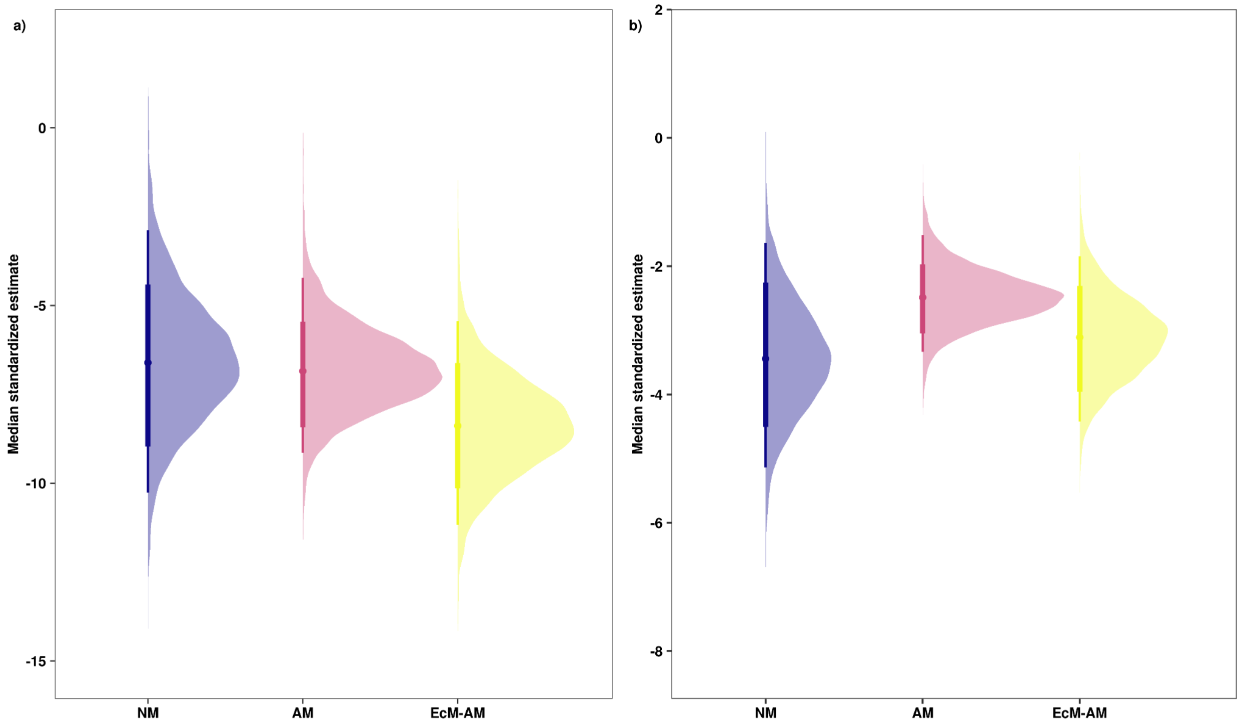


**Figure S13.** Coefficient estimates of the effect of mycorrhizal types to predict the **a**) establishment (presence/absence) and **b)** dominance (relative abundance) of alien woody species across the Hawaiian archipelago. Mycorrhizal types were assigned by plant genera; NM is no mycorrhiza, AM is arbuscular mycorrhiza, and EcM-AM is ectomycorrhiza and arbuscular mycorrhiza. Medians, 80% and 95% credible intervals of model coefficients were estimated using a phylogenetic hierarchical Bayesian with a Bernoulli distribution.

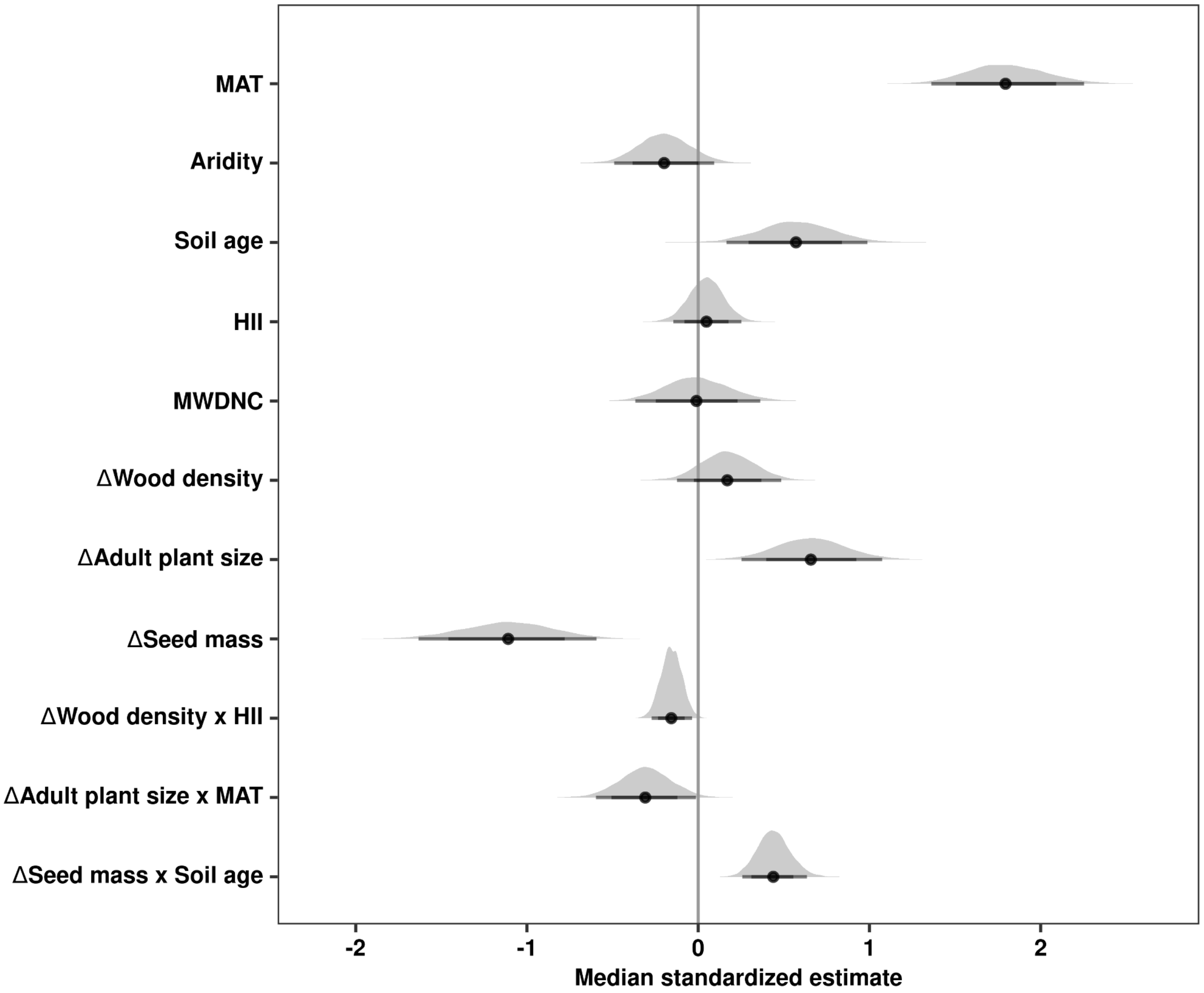


**Figure S14.** Coefficient estimates of main effects used to predict the establishment of alien woody species across the Hawaiian archipelago when excluding *M. polymorpha* from the dataset. Aridity is the ratio of mean annual potential evapotranspiration (PET) to mean annual precipitation, MAT is mean annual temperature, soil age is the estimated age of soil substrate, HII is the human influence index, MWDNC, i.e. phylogenetic distinctiveness, is the mean weighted phylogenetic distance to the native tree community, and ΔWD, Δ seed mass, and Δ max. height are the standardised differences in wood density, seed mass, and maximum height, respectively, between alien woody species and the native tree community species. Medians, 80% and 95% credible intervals of model coefficients were estimated using a phylogenetic hierarchical Bayesian with a Bernoulli distribution.

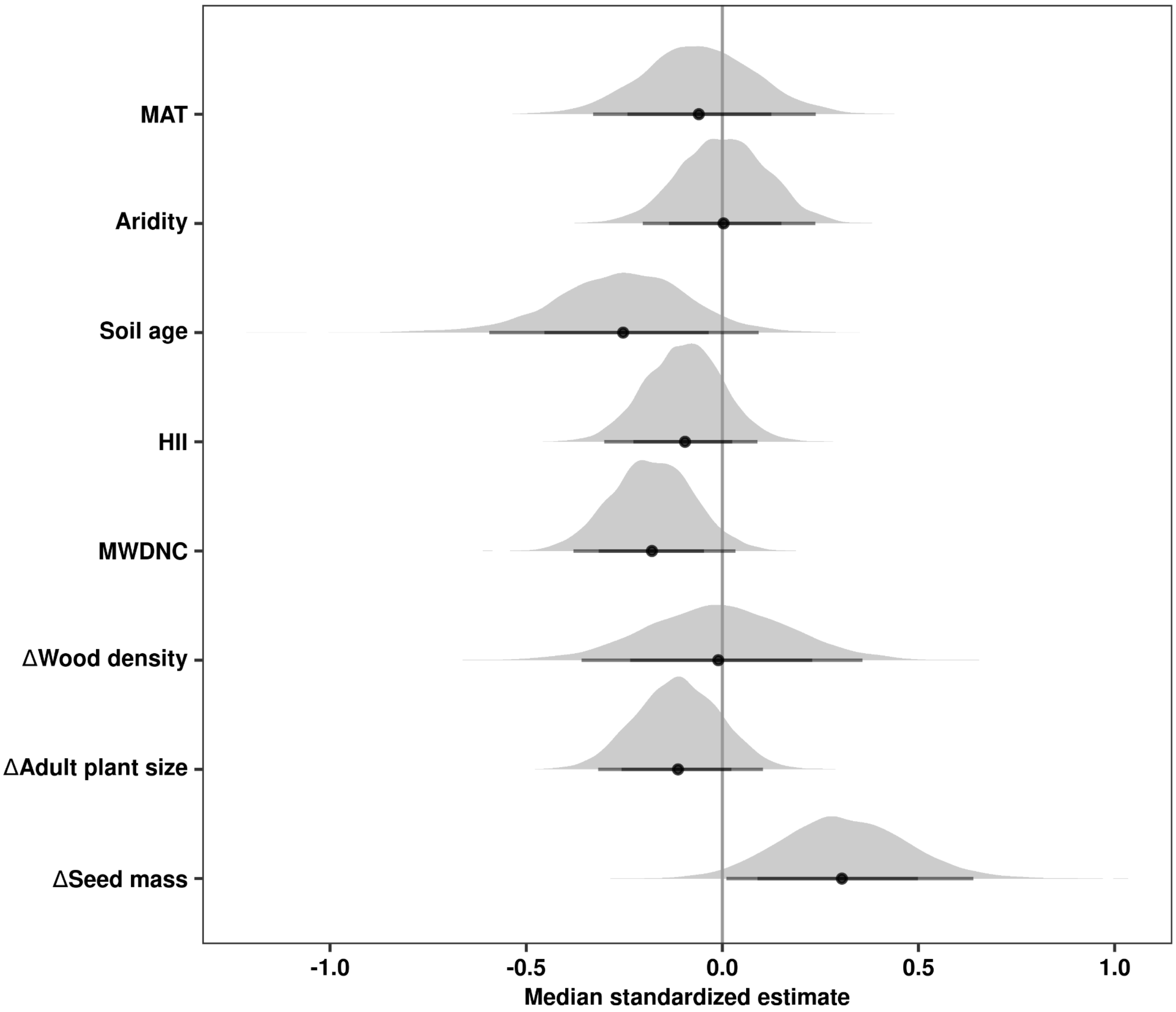


**Figure S15**. Coefficient estimates of main effects used to predict dominance of alien woody species across Hawaiian forests, when excluding *M. polymorpha* from the dataset. Aridity is the ratio of mean annual potential evapotranspiration (PET) to mean annual precipitation, MAT is mean annual temperature, soil age is the estimated age of soil substrate, HII is the human influence index, MWDNC, i.e. phylogenetic distinctiveness, is the mean weighted phylogenetic distance to the native tree community, and ΔWD, Δ seed mass, and Δ max. height are the standardised differences in wood density, seed mass, and maximum height, respectively, between alien woody species and the native tree community species. Medians, 80%, and 95% credible intervals of model coefficients were estimated using a phylogenetic hierarchical Bayesian model with a log-normal distribution.

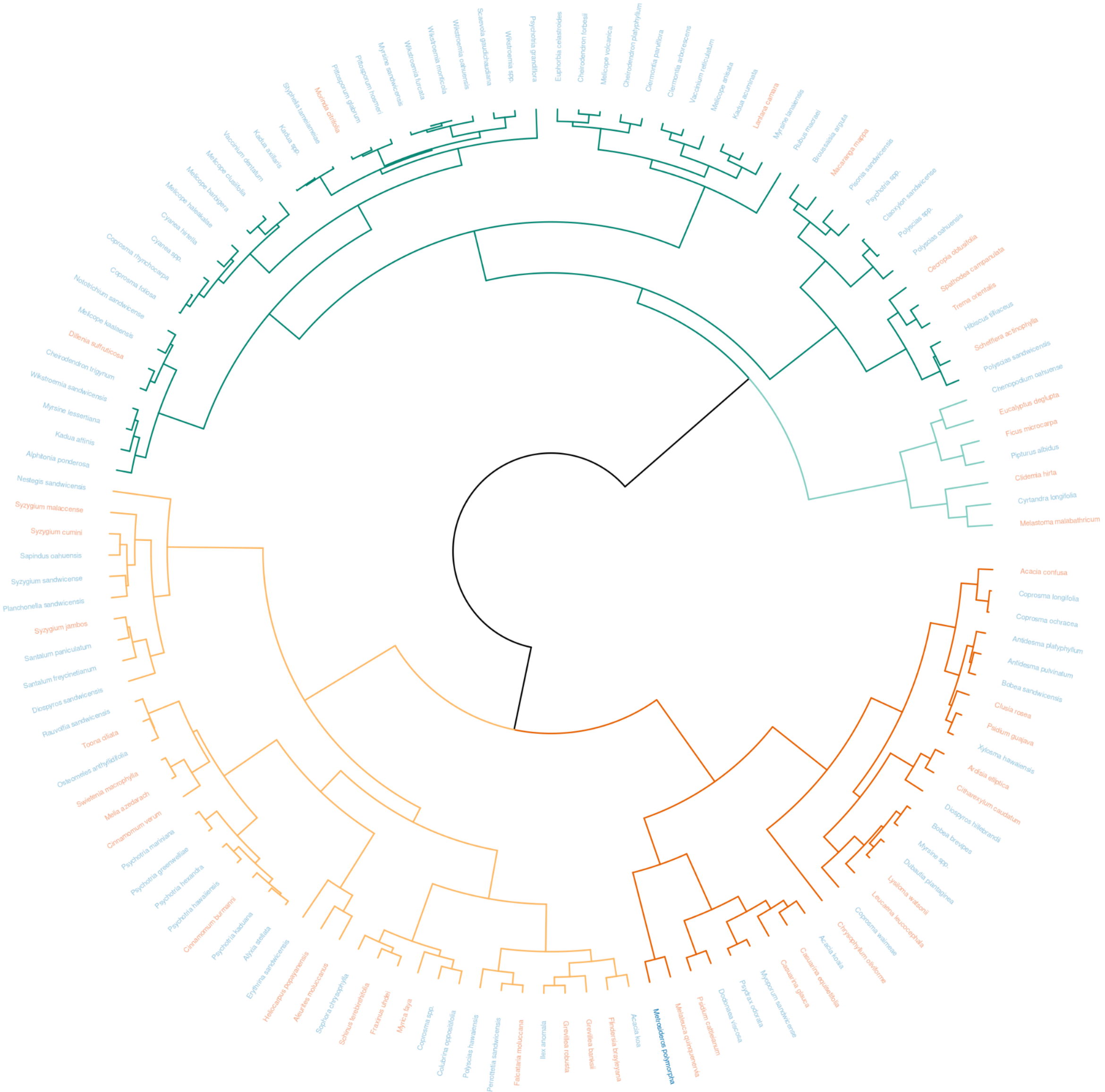

**Figure S16.** Dendrogram of native and alien woody species in Hawaiian forests clustered by functional traits (wood density, seed mass, adult plant size). Agglomerative hierarchical clustering was performed on scaled and transformed trait data using Euclidean distance and Ward's method. The most parsimonious number of clusters was determined by average silhouette width (k = 4, average silhouette width = 0.27). Branch colors correspond to clusters (group 1 = dark orange, group 2= light orange, group 3 = dark green, group 4 = light green), and label colors indicate either native (blue) or alien (orange) species. *Metrosideros polymorpha* is in bold.

**Table S1.** List of native and alien woody species present in plot database of Hawaiian forests, including scientific name, native status, wood density, adult plant size, and seed mass. When data were not available at the species level for plant functional traits, values were imputed using phylogenetic information. Imputed values are indicated with “I” .

| Scientific name | Native status | Wood density ($g\ cm^{-3}$) | Adult plant size (cm) | Seed mass (mg) |
|---|---|---|---|---|
| *Acacia confusa* | Alien | 0.73 | 21.27 | 32 |
| *Acacia koa* | Native | 0.55 | 86.25 | 102.4 |
| *Acacia koaia* | Native | 0.77 | 48.44 | 11.09 |
| *Aleurites moluccanus* | Alien | 0.38 | 37.23 | 7475.12 |
| *Alphitonia ponderosa* | Native | 0.5 | 28.93 | 36.46 (I) |
| *Alyxia stellata* | Native | 0.5 | 24.10 | 102.94 (I) |
| *Antidesma platyphyllum* | Native | 0.67 | 21.67 | 20.84 (I) |
| *Antidesma pulvinatum* | Native | 0.62 | 23.63 | 20.84 (I) |
| *Ardisia elliptica* | Alien | 0.62 | 16.97 | 84.4 (I) |
| *Bobea brevipes* | Native | 0.65 | 14.00 | 22.51 |
| *Bobea sandwicensis* | Native | 0.65 | 26.49 | 28.71 |
| *Broussaisia arguta* | Native | 0.21 | 15.80 | 69.8 |
| *Casuarina equisetifolia* | Alien | 0.81 | 37.68 | 4.18 |
| *Casuarina glauca* | Alien | 0.75 | 26.80 | 2.74 |
| *Cecropia obtusifolia* | Alien | 0.29 | 27.93 | 1.7 |
| *Cheirodendron forbesii* | Native | 0.47 | 9.65 | 10.36 |
| *Cheirodendron platyphyllum* | Native | 0.47 | 8.70 | 4.32 |

| **Scientific name** | **Native status** | **Wood density ($g\ cm^{-3}$)** | **Adult plant size (cm)** | **Seed mass (mg)** |
|---|---|---|---|---|
| *Cheirodendron trigynum* | Native | 0.47 | 28.34 | 15.09 |
| *Chenopodium oahuense* | Native | 0.5 | 16.76 | 0.68 (I) |
| *Chrysophyllum oliviforme* | Alien | 0.82 | 5.33 | 2045.14 (I) |
| *Cinnamomum burmanni* | Alien | 0.49 | 27.81 | 97.3 |
| *Cinnamomum verum* | Alien | 0.49 | 23.27 | 568.2 |
| *Citharexylum caudatum* | Alien | 0.65 | 20.76 | 134 |
| *Claoxylon sandwicense* | Native | 0.35 | 18.52 | 10 (I) |
| *Clermontia arborescens* | Native | 0.5 | 6.77 | 6.75 |
| *Clermontia parviflora* | Native | 0.5 | 8.33 | 6.98 |
| *Clidemia hirta* | Alien | 0.64 (I) | 5.00 | 0.03 |
| *Clusia rosea* | Alien | 0.63 | 18.88 | 11.07 |
| *Colubrina oppositifolia* | Native | 0.7 | 32.80 | 29.22 (I) |
| *Coprosma foliosa* | Native | 0.48 | 14.76 | 15.24 (I) |
| *Coprosma longifolia* | Native | 0.71 | 16.21 | 15.24 (I) |
| *Coprosma ochracea* | Native | 0.71 | 17.00 | 15.24 (I) |
| *Coprosma rhynchocarpa* | Native | 0.48 | 14.23 | 15.24 (I) |
| *Coprosma spp.* | Native | 0.69 (I) | 44.50 | 15.24 (I) |
| *Coprosma waimeae* | Native | 0.71 | 8.28 | 15.24 (I) |
| *Cyanea hirtella* | Native | 0.5 (I) | 12.20 | 13.54 |

| Scientific name | Native status | Wood density (g cm$^{-3}$) | Adult plant size (cm) | Seed mass (mg) |
|---|---|---|---|---|
| *Cyanea spp.* | Native | 0.5 (I) | 16.00 | 13.39 |
| *Cyrtandra longifolia* | Native | 0.5 (I) | 8.45 | 0.01 (I) |
| *Dillenia suffruticosa* | Alien | 0.45 | 28.13 | 9 |
| *Diospyros hillebrandii* | Native | 0.7 | 16.95 | 385.46 (I) |
| *Diospyros sandwicensis* | Native | 0.74 | 51.70 | 385.46 (I) |
| *Dodonaea viscosa* | Native | 0.95 | 26.42 | 6.86 |
| *Dubautia plantaginea* | Native | 0.65 | 12.00 | 25.71 |
| *Erythrina sandwicensis* | Native | 0.25 | 66.69 | 543.04 |
| *Eucalyptus deglupta* | Alien | 0.45 | 32.16 | 0.16 |
| *Euphorbia celastroides* | Native | 0.4 (I) | 10.05 | 8.13 (I) |
| *Falcataria moluccana* | Alien | 0.42 | 89.55 | 24.26 |
| *Ficus microcarpa* | Alien | 0.41 | 12.50 | 0.33 |
| *Flindersia brayleyana* | Alien | 0.48 | 81.00 | 93.98 |
| *Fraxinus uhdei* | Alien | 0.57 (I) | 39.00 | 23.98 |
| *Grevillea banksii* | Alien | 0.55 | 74.88 | 29.45 |
| *Grevillea robusta* | Alien | 0.52 | 53.60 | 16.65 |
| *Heliocarpus popayanensis* | Alien | 0.24 | 43.81 | 22300 |
| *Hibiscus tilliaceus* | Native | 0.42 (I) | 38.94 | 16 |
| *Ilex anomala* | Native | 0.48 | 59.12 | 17.71 (I) |

| Scientific name | Native status | Wood density (g cm$^{-3}$) | Adult plant size (cm) | Seed mass (mg) |
|---|---|---|---|---|
| *Kadua acuminata* | Native | 0.56 | 7.46 | 12.51 |
| *Kadua affinis* | Native | 0.56 | 23.62 | 18.26 |
| *Kadua axillaris* | Native | 0.55 | 15.66 | 15.85 |
| *Kadua spp.* | Native | 0.56 | 15.42 | 15.77 |
| *Lantana camara* | Alien | 0.6 | 6.81 | 11.05 |
| *Leucaena leucocephala* | Alien | 0.67 | 15.89 | 42.75 |
| *Lysiloma watsonii* | Alien | 0.67 (I) | 11.25 | 31.59 (I) |
| *Macaranga mappa* | Alien | 0.27 | 18.36 | 24.94 (I) |
| *Melaleuca quinquenervia* | Alien | 0.65 | 71.46 | 0.28 |
| *Melastoma malabathricum* | Alien | 0.44 | 11.40 | 0.06 |
| *Melia azedarach* | Alien | 0.43 | 34.72 | 779.84 |
| *Melicope anisata* | Native | 0.45 | 6.30 | 11.94 (I) |
| *Melicope barbigera* | Native | 0.45 | 18.80 | 11.94 (I) |
| *Melicope clusiifolia* | Native | 0.45 | 16.56 | 11.94 (I) |
| *Melicope haleakalae* | Native | 0.45 | 12.05 | 11.94 (I) |
| *Melicope kaalaensis* | Native | 0.45 | 22.00 | 11.94 (I) |
| *Melicope volcanica* | Native | 0.45 | 9.34 | 11.94 (I) |
| *Metrosideros polymorpha* | Native | 0.69 | 82.82 | 0.04 |
| *Morinda citrifolia* | Alien | 0.56 | 11.30 | 15 |

| Scientific name | Native status | Wood density (g cm$^{-3}$) | Adult plant size (cm) | Seed mass (mg) |
|---|---|---|---|---|
| *Myoporum sandwicense* | Native | 0.88 | 39.15 | 23.68 (I) |
| *Myrica faya* | Alien | 0.66 | 50.00 | 4.7 |
| *Myrsine lanaiensis* | Native | 0.53 | 6.50 | 25.03 (I) |
| *Myrsine lessertiana* | Native | 0.53 | 19.74 | 25.03 (I) |
| *Myrsine sandwicensis* | Native | 0.53 | 13.22 | 25.03 (I) |
| *Myrsine spp.* | Native | 0.63 (I) | 13.52 | 25.03 (I) |
| *Nestegis sandwicensis* | Native | 0.66 | 129.15 | 550 (I) |
| *Nototrichium sandwicense* | Native | 0.48 | 23.88 | 21.71 |
| *Osteomeles anthyllidifolia* | Native | 0.45 | 42.15 | 133.21 |
| *Perrottetia sandwicensis* | Native | 0.41 | 73.47 | 26.82 |
| *Pipturus albidus* | Native | 0.3 | 12.57 | 0.45 (I) |
| *Pisonia sandwicensis* | Native | 0.3 | 21.93 | 87.51 (I) |
| *Pittosporum glabrum* | Native | 0.57 | 10.60 | 15.85 (I) |
| *Pittosporum hosmeri* | Native | 0.57 | 9.79 | 15.85 (I) |
| *Planchonella sandwicensis* | Native | 0.69 (I) | 53.77 | 1606.02 |
| *Polyscias hawaiensis* | Native | 0.35 (I) | 62.10 | 10.99 (I) |
| *Polyscias oahuensis* | Native | 0.35 (I) | 24.56 | 10.99 (I) |
| *Polyscias sandwicensis* | Native | 0.35 (I) | 39.63 | 10.99 (I) |
| *Polyscias spp.* | Native | 0.35 (I) | 17.50 | 10.99 (I) |
| *Psidium cattleianum* | Alien | 0.98 | 20.12 | 15.46 |

| Scientific name | Native status | Wood density (g cm$^{-3}$) | Adult plant size (cm) | Seed mass (mg) |
|---|---|---|---|---|
| *Psidium guajava* | Alien | 0.66 | 23.46 | 9.77 |
| *Psychotria grandiflora* | Native | 0.52 | 9.34 | 122.56 (I) |
| *Psychotria greenwelliae* | Native | 0.52 | 31.18 | 122.56 (I) |
| *Psychotria hawaiiensis* | Native | 0.54 | 22.82 | 122.56 (I) |
| *Psychotria hexandra* | Native | 0.52 | 20.44 | 122.56 (I) |
| *Psychotria kaduana* | Native | 0.52 | 26.04 | 122.56 (I) |
| *Psychotria mariniana* | Native | 0.52 | 34.99 | 122.56 (I) |
| *Psychotria spp.* | Native | 0.34 (I) | 15.00 | 122.56 (I) |
| *Psydrax odorata* | Native | 0.87 | 35.70 | 13.07 |
| *Rauvolfia sandwicensis* | Native | 0.46 | 45.66 | 68.36 (I) |
| *Rubus macraei* | Native | 0.31 (I) | 5.10 | 3.99 (I) |
| *Santalum freycinetianum* | Native | 0.75 | 35.74 | 922.5 (I) |
| *Santalum paniculatum* | Native | 0.75 | 24.35 | 922.5 (I) |
| *Sapindus oahuensis* | Native | 0.63 (I) | 36.09 | 943.59 (I) |
| *Scaevola gaudichaudiana* | Native | 0.5 | 9.70 | 38.34 (I) |
| *Schefflera actinophylla* | Alien | 0.41 | 31.91 | 7.24 |
| *Schinus terebinthifolia* | Alien | 0.62 (I) | 34.48 | 18.11 |
| *Sophora chrysophylla* | Native | 0.6 | 50.68 | 39.99 |
| *Spathodea campanulata* | Alien | 0.34 | 45.59 | 1.99 |
| *Styphelia tameiameiae* | Native | 0.56 | 15.00 | 14.61 (I) |

| **Scientific name** | **Native status** | **Wood density (g cm$^{-3}$)** | **Adult plant size (cm)** | **Seed mass (mg)** |
|---|---|---|---|---|
| *Swietenia macrophylla* | Alien | 0.42 | 34.33 | 542.02 |
| *Syzygium cumini* | Alien | 0.65 | 47.85 | 689.04 |
| *Syzygium jambos* | Alien | 0.7 | 28.28 | 1463.03 |
| *Syzygium malaccense* | Alien | 0.56 | 31.22 | 2465.34 |
| *Syzygium sandwicense* | Native | 0.63 | 40.94 | 3523.91 (I) |
| *Toona ciliata* | Alien | 0.42 | 41.73 | 80.28 |
| *Trema orientalis* | Alien | 0.35 | 40.64 | 4.4 |
| *Vaccinium dentatum* | Native | 0.51 (I) | 17.34 | 5.08 (I) |
| *Vaccinium reticulatum* | Native | 0.51 (I) | 5.78 | 5.08 (I) |
| *Wikstroemia furcata* | Native | 0.52 | 13.90 | 26.23 (I) |
| *Wikstroemia monticola* | Native | 0.52 | 13.85 | 26.23 (I) |
| *Wikstroemia oahuensis* | Native | 0.52 | 12.11 | 26.23 (I) |
| *Wikstroemia sandwicensis* | Native | 0.52 | 22.90 | 26.23 (I) |
| *Wikstroemia spp.* | Native | 0.52 (I) | 10.40 | 26.23 (I) |
| *Xylosma hawaiensis* | Native | 0.65 (I) | 27.58 | 9.12 (I) |

**Table S2.** Summary of taxonomic resolution of trait data and the percentage of trait records imputed using phylogenetic information.

| **Trait** | **% records at species level** | **% records at genus level** | **% records imputed** |
|---|---|---|---|
| Adult plant size | 99% | 0 | 1% |
| Seed mass | 36% | 50% | 13% |
| Wood density | 77% | 18% | 5% |

**Table S3**. Model convergence and explained variation of phylogenetic hierarchical models that examine relative species richness and relative abundance of alien woody plant communities in Hawaiian forests. Model convergence was evaluated by estimating '*Rhat*', where values greater than 1.01 indicate that models have failed to converge; Bayesian $r^2$ represents an estimate of the proportion of variation explained for new data.

| **Indicator of biological invasion** | **Rhat** | **Bayesian $R^2$** |
|---|---|---|
| Rel. species richness | 1.00056 (1.00025-1.00087) | 0.35 (0.26-0.44) |
| Rel. abundance | 1.0004 (0.99999-1.00081) | 0.18 (0.10-0.26) |

**Table S4.** Model convergence and explained variation of phylogenetic hierarchical models that examine establishment (presence/absence) and dominance (relative abundance) of alien woody species in Hawaiian forests. Model convergence was evaluated by estimating '*Rhat*', where values greater than 1.01 indicate that models have failed to converge; Bayesian $r^2$ represents an estimate of the proportion of variation explained for new data.

| **Invasion stage** | **Rhat** | **Bayesian $r^2$** |
|---|---|---|
| Establishment | 1.00005 (1-1.0001) | 0.29 (0.25-0.32) |
| Dominance | 1.00003 (0.99996-1.0001) | 0.47 (0.35-0.57) |

**Table S5**. Summary of a phylogenetic hierarchical model that examines establishment (presence/absence) of alien woody species in Hawaiian forests. Estimates are the median of fixed effects; numbers in parentheses are 95% uncertainty intervals. Effective sample size represents the number of independent draws that contain the same amount of information as the dependent sample obtained by the MCMC algorithm.

| Fixed effect | Estimate | Effective sample size |
|---|---|---|
| Intercept | -7.01 (-9.18, -4.62) | 2010.540 |
| Aridity | 0.24 (-0.08, 0.59) | 4272.429 |
| MAT | 2.00 (1.58, 2.46) | 4407.406 |
| HII | 0.18 (-0.01, 0.38) | 5655.417 |
| Soil age | 0.47 (0.12, 0.82) | 5345.062 |
| MWDNC | -0.08 (-0.37, 0.22) | 6079.616 |
| Δ Wood density | 0.06 (-0.33, 0.45) | 3550.174 |
| Δ Adult plant size | 0.55 (0.09, 1.00) | 3047.736 |
| Δ Seed mass | -0.09 (-0.48, 0.29) | 3871.578 |
| MWDNC x MAT | -0.42 (-0.66, -0.17) | 4953.021 |
| MWDNC x Aridity | 0.33 (0.13, 0.54) | 5242.461 |
| Δ Wood density x HII | -0.21 (-0.32, -0.1) | 8199.418 |
| Δ Wood density x Soil age | 0.27 (0.16, 0.38) | 7854.987 |
| Δ Adult plant size x Aridity | 0.45 (0.17, 0.71) | 6709.894 |
| Δ Adult plant size x MAT | -0.68 (-1.01, -0.35) | 4404.426 |

| Fixed effect | Estimate | Effective sample size |
|---|---|---|
| Δ Seed mass x MAT | -0.54 (-0.87, -0.19) | 4770.548 |
| Δ Seed mass x Aridity | 0.34 (0.06, 0.62) | 5731.063 |
| Δ Seed mass x Soil age | 0.35 (0.2, 0.5) | 5331.824 |

**Table S6**. Summary of a phylogenetic hierarchical model that examines dominance (relative abundance) of alien woody species in Hawaiian forests. Estimates are the median of fixed effects; numbers in parentheses are 95% uncertainty intervals. Effective sample size represents the number of independent draws that contain the same amount of information as the dependent sample obtained by the MCMC algorithm.

| Fixed effect | Estimate | Effective sample size |
|---|---|---|
| Intercept | -2.51 (-3.32, -1.63) | 4391.599 |
| Aridity | -0.05 (-0.24, 0.13) | 7699.316 |
| MAT | -0.04 (-0.29, 0.21) | 6247.028 |
| HII | -0.08 (-0.25, 0.09) | 8321.256 |
| Soil age | -0.11 (-0.41, 0.16) | 6103.228 |
| MWDNC | -0.23 (-0.48, 0.01) | 7711.772 |
| Δ Wood density | -0.1 (-0.48, 0.29) | 6048.186 |
| Δ Adult plant size | -0.09 (-0.31, 0.13) | 5779.955 |
| Δ Seed mass | 0.06 (-0.13, 0.24) | 5987.837 |
